# Comparison of three tiled amplicon sequencing approaches for SARS-CoV-2 variant detection from wastewater

**DOI:** 10.1101/2024.06.16.599198

**Authors:** Megan E. J. Lott, Amanda H. Sullivan, Leah M. Lariscy, William A. Norfolk, Katie C. Dillon, Megan S. Beaudry, Travis C. Glenn, Erin K. Lipp

**Affiliations:** Department of Environmental Health Science, University of Georgia, Athens GA 30602 USA; Institute of Bioinformatics, University of Georgia, Athens GA 30602 USA

## Abstract

During the COVID-19 pandemic, the detection and sequencing of SARS-CoV-2 from wastewater proved to be a valuable tool in assessing trends at the community level. Several whole genome enrichment methods have been proposed for sequencing SARS-CoV-2 from the mixed wastewater community, but there is little consensus on the most appropriate sequencing methods for variant detection or abundance estimations. Few studies have elucidated the errors associated with these methods or have established minimum sequencing requirements for correct interpretation of the results. To address these needs, we systematically assessed the efficacy of three tiled amplicon enrichment methods (Freed/Midnight, ARTIC V4, NEB VarSkip) for whole genome sequencing of SARS-CoV-2 variants using mock wastewater communities with variants at known proportions. We found the ARTIC V4 approach yielded the most accurate results for variant identification and variant abundance estimation, followed by the NEB VarSkip approach. Conversely, the NEB VarSkip method obtained the highest genomic coverage, with the ARTIC V4 method achieving the second highest coverage. Finally, we determined that the Freed/Midnight library preparation methods are not well-suited for use with short read sequencing. Based on the present results, the ARTIC V4 workflow appears to be the most robust and cost-effective approach for monitoring circulating SARS-CoV-2 variants with wastewater surveillance.

**IMPORTANCE:** This work is informative for practitioners of wastewater-based epidemiology. Here, we detail a systematic comparison of three tiled amplicon sequencing approaches for enrichment of SARS-CoV-2 variants from wastewater. Using mock communities of known variant composition, we validate the analysis methods previously published by Baaijens et al. in Genome Biology (2022) for estimating variant abundance from wastewater using an RNAseq pipeline, kallisto. We provide recommendations for minimum sequencing requirements for accurate abundance estimates of SARS-CoV-2 variants in wastewater. The sequences generated from the mock communities have been uploaded to NCBI’s Sequence Read Archive and will be useful to other practitioners seeking to validate their sequencing methods or bioinformatic pipelines.

## INTRODUCTION

Wastewater-based epidemiology (WBE) is a robust approach for disease surveillance in which wastewater samples are viewed as a pooled collective sample that captures a snapshot of the community’s health, without the need for extensive clinical testing. In response to the COVID-19 pandemic, wastewater surveillance for SARS-CoV-2 has been adopted globally to monitor epidemic progression at local scales (Daleiden et al., 2022; Kirby et al., 2021). Viral titers of SARS-CoV-2 in wastewater have been demonstrated to correlate with reported clinical cases of COVID-19 (Larsen & Wigginton, 2020; Medema, Heijnen, Elsinga, Italiaander, & Brouwer, 2020; Polo et al., 2020). In communities where clinical testing is limited, wastewater surveillance has effectively predicted local outbreaks or surges, with lead times up to several days ahead of clinical testing (Bibby, Bivins, Wu, & North, 2021; Olesen, Imakaev, & Duvallet, 2021; Zhu et al., 2021).

WBE has been paired with molecular epidemiology to monitor circulating and emerging variants of SARS-CoV-2. With the continued emergence and circulation of novel SARS-CoV-2 lineages, variant-specific detection assays have been employed in wastewater surveillance to monitor introduction events of novel variants into local communities (Kirby et al., 2022; Yu et al., 2021). By sequencing SARS-CoV-2 genomes in wastewater, several groups have predicted SARS-CoV-2 variant abundance that correlates with clinical trends, and others have identified emerging or cryptic lineages in wastewater that were not captured by clinical sequencing (Baaijens et al., 2021; Crits-Christoph et al., 2021; Fontenele et al., 2021; Karthikeyan et al., 2022; Nemudryi et al., 2020; Rouchka et al., 2021; Schumann et al., 2022; Smyth et al., 2022). In fact, Baaijens et al. determined that different variants of SARS-CoV-2 could be estimated with a fair amount of accuracy with the RNA-seq program kallisto and a reference database containing SARS-CoV-2 variant genomes.

Several sequencing methods have been described for SARS-CoV-2 variant monitoring in wastewater (Barbé et al., 2022; Lin et al., 2021; Ni et al., 2021). Most often, these approaches have been adapted from workflows that were originally developed for short read sequencing, such as those described by the ARTIC network (Quick, 2020), New England BioLabs (Grim, 2022), and Freed et al. (2021). The Freed/Midnight workflow generates long tiled amplicons of approximately 1,200 bp, whereas the NEB VarSkip and ARTIC V4 workflows generate amplicons of approximately 560 bp and 400 bp, respectively. There is no clear consensus on which of these methods is the most appropriate for sequencing SARS-CoV-2 from wastewater, a notoriously difficult sample matrix. Viral genomes in wastewater are low in titer and are often heavily degraded (Wurtzer et al., 2021). Genomic enrichment is often required prior to sequencing, however tiled amplicon assays are susceptible to inhibition, off-target amplification, and replication errors (Lin et al., 2021). While clinical samples are comprised of a single variant, wastewater samples are comprised of multiple strains (and potentially multiple variants), pooled from an entire community, making it challenging to detect low-frequency mutations. Despite these challenges, few studies have fully elucidated the errors associated with amplicon sequencing from wastewater or systematically compared methods to assess biases introduced at the amplicon and sequencing levels.

In this study, we evaluated the use of three tiled amplicon enrichment methods for whole genome sequencing of SARS-CoV-2 from wastewater, the Freed/Midnight workflow, the NEB VarSkip workflow and the ARTIC V4 workflow. We will use the method created by Baaijeens et al. to identify variants and estimate their abundancies. By sequencing mock wastewater communities composed of variants at known proportions, we aimed to systematically compare these currently available and widely used methods, while assessing the challenges and errors associated with these sequencing approaches.

## RESULTS

### Sequencing Statistics

Fifteen mock communities, three positive controls, and two negative controls were enriched with Midnight, VarSkip and ARTIC V4 tiled amplicon primer sets. From these, 60 libraries were sequenced with PE150 reads, 60 libraries were sequenced with PE250 reads, and 17 libraries were sequenced with PE300 reads. Across all libraries, 40,635,190 raw PE150 reads, 560,282 raw PE250 reads, and 484,453 PE300 reads were generated using Illumina chemistry (Table 1). After quality filtering, 17,083,262 PE150 reads (42%), 483,802 PE250 reads (86%), and 443,199 PE300 reads (92%) were retained.

**Table 1.** Sequence Summary Table. Summary of sequences obtained from whole genome amplification of SARS-CoV-2 from wastewater mock communities (all variant combinations and final concentrations, combined), using three tiled amplicon approaches. Assignment, breadth of coverage, and depth are all relative to filtered reads.

| Primer Set | Read Length | No. Samples | Raw Reads<br>Median (Range) | Filtered Reads<br>Median (Range) | Read Retention (%)<br>Median (Range) | Reads Assigned in Kallisto<br>Median (Range) | Read Assignment (%)<br>Median (Range) | Genome Breadth of Coverage (%)<br>Median (Range) | Depth<br>Median (Range) | Genome Coverage >20X Depth (%)<br>Median (Range) |
| --- | --- | --- | --- | --- | --- | --- | --- | --- | --- | --- |
| Midnight | PE150 | 15 | 925,520<br>(149,940-1,474,862) | 377,458<br>(47,248-1,474,862) | 44<br>(4-45) | 329,005<br>(23,648-535,350) | 89<br>(46-96) | 82<br>(75-89) | 480<br>(15-1,879) | 71<br>(45-82) |
| Midnight | PE250 | 15 | 13,175<br>(1,926-18,968) | 11,840<br>(1,769-18,968) | 90<br>(84-92) | 9,672<br>(1,652-16,349) | 90<br>(52-96) | 70<br>(45-82) | 18<br>(<1-46) | 50<br>(14-60) |
| ARTIC V4 | PE150 | 15 | 289,996<br>(7,544-544,970) | 118,348<br>(3,059-544,970) | 41<br>(40-41) | 106,940<br>(1,035-189,708) | 88<br>(34-97) | 94<br>(66-97) | 181<br>(3-516) | 72<br>(15-87) |
| ARTIC V4 | PE250 | 15 | 2,030<br>(90-4,686) | 1,813<br>(73-4,686) | 93<br>(81-96) | 1,527<br>(23-3,598) | 86<br>(32-95) | 84<br>(16-95) | 17<br>(<1-30) | 46<br>(0-64) |
| VarSkip | PE150 | 15 | 448,658<br>(168,500-3,882,174) | 198,844<br>(70,570-388,2174) | 43<br>(42-45) | 191,692<br>(6,9421-1,694,581) | 98<br>(96-98) | 97<br>(95-99) | 854<br>(356-5,276) | 94<br>(90-96) |

The 45 mock community libraries sequenced with PE150 chemistry generated between 10^5^ and 10^6^ reads each; the 45 mock community libraries sequenced with PE250 chemistry generated between 10^3^ and 10^4^ reads each, and the 15 mock community libraries sequenced with PE300 chemistry generated approximately 10^4^ reads each (Table 1). After quality filtering, read retention for the mock communities ranged between 4% and 96% (Table 1). Library preparation with ARTIC V4 tiled amplicons and subsequent sequencing with PE250 chemistry resulted in the greatest median read retention 93% (81% - 96%) (Table 1). Sequencing with PE150 chemistry reduced retention of V4 amplicon reads significantly (Dunn’s pairwise test, *p*_Holm-adj_ < 0.001). Approximately 90% (Table 1) of raw reads were retained from Midnight amplicons when sequenced with PE250 chemistry, but median retention was significantly reduced to 44% (4% - 45%) when Midnight amplicons were sequenced with PE150 chemistry (Dunn’s pairwise test, *p*_Holm-adj_ < 0.001). Approximately 75% (35% - 93%) of reads were retained from the VarSkip libraries when sequenced with PE300 chemistry, comparable to median number of reads were retained when sequenced with PE250 chemistry, 81% (75% - 91%). Significantly fewer reads, 43% (42% - 45%), were retained when VarSkip libraries were sequenced with PE150 than with PE300 chemistry (Dunn’s pairwise test, *p*_Holm-adj_ < 0.001).

### Genomic Enrichment

Overall, the positive controls obtained high breadth of genomic coverage and high depth of genomic coverage. The Wuhan Twist Control had between 85% and 96% breadth of coverage, with median depth of coverage ranging from 9.6X to 1774X across the three library preparation methods (Supplemental Figure 1). The genomic breadth coverage of the heat-inactivated SARS-CoV-2 control carried in PBS when using three tiled amplicon preparation methods ranged from approximately 67% to 99%, while the median depth of coverage ranged from 63X to 1632X (Supplemental Figure 2). For the heat-inactivated SARS-CoV-2 control spiked into wastewater the genomic breadth of coverage and the genomic depth of coverage obtained using the three library preparation methods was generally lower, with genomic breadth of coverage ranging from 5% to 84% and median depth of coverage ranging from 0.14% to 1824 (Supplemental Figure 3). This suggests that the library prep methods are inhibited by the addition of wastewater, though the V4 library prep method still obtained fair breadth of coverage (63% with PE250 reads and 84% with PE150 reads) with the wastewater present.

Sequencing reads from mock community libraries covered between 16% and 99% of the SARS-CoV-2 Wuhan reference genome (Table 1, Supplemental Figures 4, 5 & 6). The sequencing depth ranged from less than 1X to more than 5,276X per sample (Table 1).

Approximately 74% (78 / 105) of the mock community samples were sequenced with a median depth greater than 20X, and 48% (50 / 105) of the samples were sequenced with a median depth greater than 100X (Supplementary Data). Few libraries (22%, 23 / 105) resulted in at least 20X depth over ≥ 90% of the genome. Variations in sequencing depth, genomic coverage, and genome coverage >20X were not attributed to RNA template concentration nor mock community composition. Instead, the efficiency and evenness of genomic enrichment varied between library preparation methods and sequencing approaches.

The median genomic coverage from all VarSkip libraries was 96% (77% - 99%) across a range of sequencing effort (Figure 1). At the highest sequencing effort, at least 10^7^ bases, genomic coverage with at least 20X depth was 96% (77% - 99%) for VarSkip libraries (Figure 1). When sequenced with at least 10^7^ bases, median genome coverage was 94% (77% – 97%) for ARTIC V4 libraries, and median genome coverage with at least 20X depth was 75% (48%-87%, Figure 1). Midnight libraries sequenced with at least 10^7^ bases resulted in 82% (75% - 89%) genome coverage and 71% (45% - 82%) genome coverage with at least 20X depth (Figure 1).

**Figure 1.**
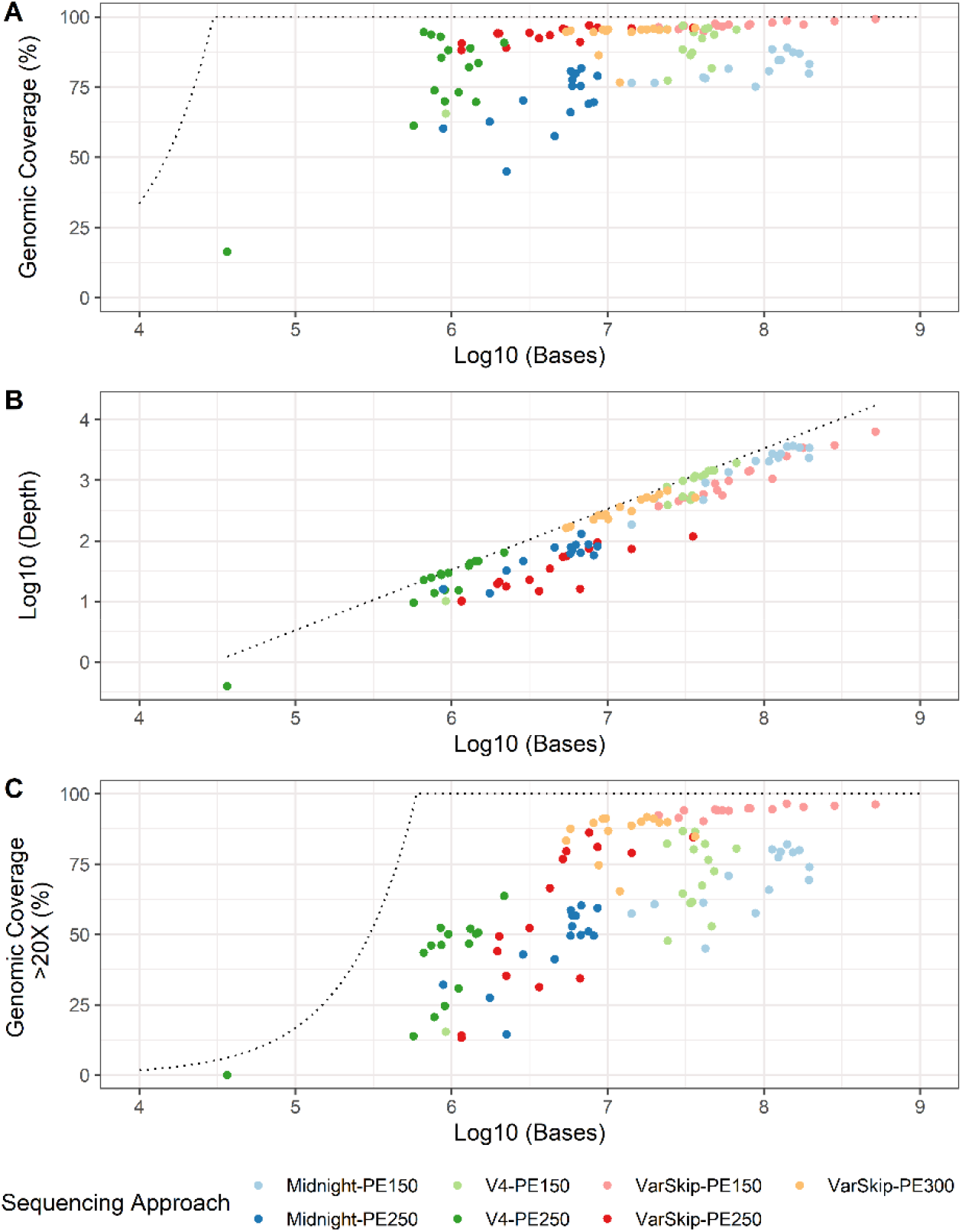
Evaluation of three tiled amplicon whole-genome enrichment methods for sequencing SARS-CoV-2 genomic RNA from wastewater mock communities. Fifteen wastewater mock communities, spiked with synthetic SARS-CoV-2 genomic RNA, were prepared as tiled amplicon libraries using Freed/Midnight, ARTIC V4, and NEB VarSkip primer schemes. Libraries were sequenced with PE250 and PE150 reads, with approximately 10^3^ and 10^4^ reads, respectively. (A) Genomic coverage from each library, as a function of sequencing effort (N = 105). The reference line represents the expected genomic coverage for the given level of sequencing effort. (B) Sequencing depth of each library, as a function of sequencing effort (N = 105). The reference line represents the expected sequencing depth for the given level of sequencing effort. (C) Sequencing depth of each library, as a function of sequencing effort (N = 105). The reference line represents the expected genomic coverage with >20X sequencing depth for the given level of sequencing effort.

### Variant Assignment with kallisto

Of the filtered sequencing reads, 32-98% were assigned to a reference variant of SARS CoV-2 by kallisto (Supplementary Data). Among the library preparation methods, VarSkip amplicons resulted in the greatest proportion of reads assigned in kallisto (Dunn’s pairwise tests, *p*_Holm-adj_ < 0.001). There was no significant difference in the proportion of assigned reads between V4 and Midnight libraries (Dunn’s pairwise test, *p*_Holm-adj_ = 0.36). Sequencing VarSkip amplicons with either PE150 or PE250 reads resulted in significantly greater proportion of assigned reads than when sequencing with PE300 reads (Dunn’s pairwise tests, *p*_Holm-adj_ < 0.001). There was no significant difference in the proportion of assigned reads when sequencing V4 amplicons with PE150 reads or with PE250 reads (Dunn’s pairwise test, *p*_Holm-adj_ = 1.0). Similarly, there was no difference in the proportion of assigned reads between Midnight amplicons sequenced with PE150 or PE250 reads (Dunn’s pairwise test, *p*_Holm-adj_ = 1.0).

Four SARS-CoV-2 variants (Wuhan, Alpha, Beta, and Delta) were spiked into each mock community at different proportions (3%, 14%, 25%, or 55%). Using a pipeline containing the program kallisto, the parent Wuhan lineage, as well as the Alpha, Beta, and Delta variants were detected in 100% of the mock community libraries (105 / 105, Supplemental Table 1). Reads were also assigned to variants of SARS-CoV-2 present in the reference database, but not spiked into the mock communities, including Epsilon, Eta, Gamma, Iota, Kappa, Mu, Omicron BA.1, Omicron BA.2, Omicron BA.4, Omicron BA.5, and Zeta. We believe these misassignments are due to low quality and short reads producing noise, as the majority of these variants had not emerged at the time these libraries were created. Reads assigned to these off-target variants are binned together as “Other” in Figure 2 but defined in detail in Supplementary Data. The most prevalent off-target assignment was Mu, called in 90% (95 / 105) of the mock community libraries, whereas the least abundant off-target assignment was Eta, called in 47% (49 / 105) of the libraries (Supplementary Data).

**Figure 2.**
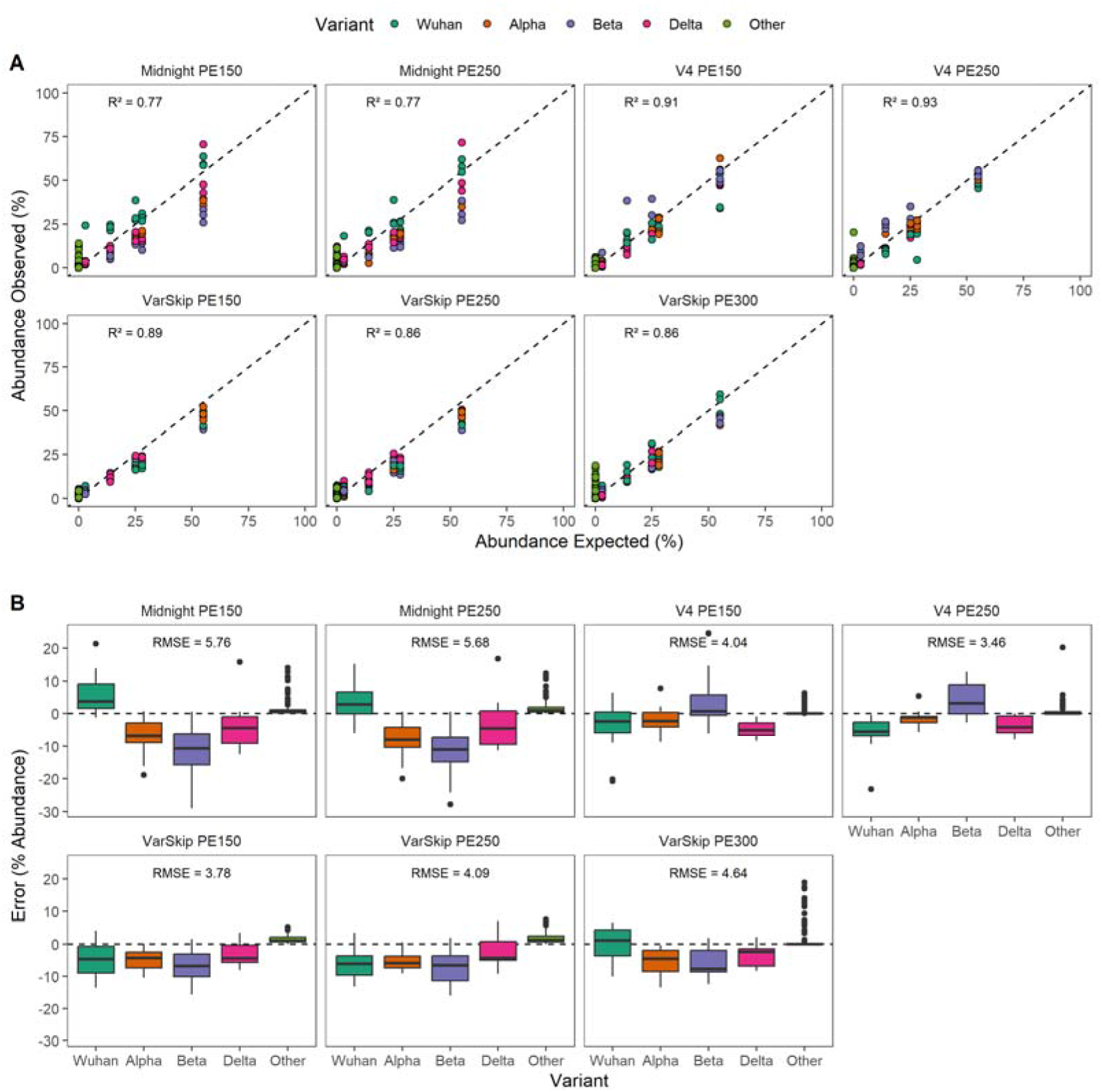
Abundance estimates of SARS-CoV-2 variants from wastewater mock communities prepared with three whole-genome tiled amplicon methods and their errors. Fifteen individual wastewater mock communities were enriched as Freed/Midnight, ARTIC V4, and NEB VarSkip libraries and sequenced with Illumina PE250 and PE150 chemistry. Sequencing reads were assessed using the kallisto workflow to estimate abundance of SARS-CoV-2 variants (Wuhan, Alpha, Beta, Delta), which were spiked into mock communities at known proportions (3%, 14%, 25%, 28%, or 55%). Reads assigned to off-target variants (Epsilon, Eta, Gamma, Iota, Kappa, Mu, Omicron and Zeta) were binned together as “Other” with an expected abundance of 0%. (A) The expected abundance of each variant was compared to observed abundance, as reported by kallisto. Accuracy of the abundance prediction was assessed as the R^2^ value of the one-to-one model between expected abundance and observed abundance of the SARS-CoV-2 variants in each sample, represented by the dashed line. (B) The prediction error associated with each SARS-CoV-2 variant expressed in RMSE. The prediction error is equivalent to the difference between the abundance reported by kallisto and the abundance expected for each variant.

### Variant Abundance

Variant assignments in kallisto were used to calculate abundance estimates for the four variants of SARS-CoV-2 (Wuhan, Alpha, Beta, Delta) spiked into each mock community sample at known proportions (Supplemental Figure 7). Across wastewater mock communities prepared with Midnight amplicons, the relative abundance of the Wuhan parent lineage was typically over-estimated, whereas the relative abundances of the Alpha, Beta, and Delta variants were typically underestimated (Figure 1). Across mock communities prepared with ARTIC V4 amplicons, the relative abundances of the Wuhan parent lineage, the Alpha, and the Delta lineages were typically under-estimated, while the Beta lineage was typically over- estimated. When prepared with VarSkip amplicons, the relative abundance of the Wuhan lineage was under-estimated, except when sequenced with PE300 reads. The relative abundances of the Alpha, Beta, and Delta lineages were typically underestimated in the VarSkip libraries, regardless of sequencing approach. Across all mock communities, the relative abundance of off-target variants ranged from 2% to 26%.

The accuracy of variant abundance prediction was assessed as the R^2^ value of the one-to-one model between expected abundance and observed abundance of the SARS-CoV-2 variants in each sample, and as the root mean squared error (RMSE) between abundance expected and abundance obtained from the data. The single variant positive controls created using the three tiled amplicon approaches have R^2^ values between -7.79 to 0.99 and RMSE values between 0.25 and 28.32 (Supplemental Figure 8-10). Within the positive controls the V4 library prep methods yielded better R^2^ and RMSE values than the VarSkip and Midnight library preparation methods. Across individual mock communities, R^2^ values ranged from 0.02 to 0.99 and RMSE values ranged from 1.13 to 8.93 (Supplemental Figures 11-13).

Based on RMSE and R^2^ values, ARTIC V4 libraries yielded the most accurate estimates of variant abundance, whereas abundance estimates from Midnight libraries were least accurate (Figure 2). The RMSE and R^2^ values were significantly greater for ARTIC V4 libraries than for Midnight libraries (Dunn’s pairwise tests, *p*_Holm-adj_ < 0.01), but not significantly different between Midnight and VarSkip libraries (Dunn’s pairwise tests, *p*_Holm-adj_ > 0.05). The R^2^ values of the ARTIC V4 libraries were not significantly different than the R^2^ values of the VarSkip libraries (Dunn’s pairwise tests, *p*_Holm-adj_ > 0.05). The RMSE values of the ARITC V4 libraries sequenced with PE250 chemistry were significantly lower than the RMSE values of VarSkip libraries sequenced with PE300 chemistry (Dunn’s pairwise tests, *p*_Holm-adj_ = 0.03), but otherwise the RMSE values between the two library methods were comparable (Dunn’s pairwise tests, *p*_Holm-adj_ > 0.05).

Neither the RMSE nor the R^2^ values were significantly different between ARTIC V4 libraries sequenced with PE150 reads and those sequenced with PE250 reads (Dunn’s pairwise test, *p*_Holm-adj_ = 1.0). Accuracy was not significantly different between VarSkip libraries sequenced with PE150, PE250, or PE300 reads, nor was the accuracy of Midnight libraries when sequenced with PE150 and PE250 reads (Dunn’s pairwise test, *p*_Holm-adj_ > 0.05, Figure 2).

When examined across all libraries, the RMSE estimates of accuracy were significantly, inversely correlated with genomic coverage (Spearman’s, Rho = -0.3, *p <* 0.01), but not with the number of sequencing reads, the median sequencing depth, nor the genome coverage > 20X (Supplemental Figure 14-15). There was no strong or significant correlation between R^2^ and the number of sequencing reads, between R^2^ and genomic coverage, between R^2^ and sequencing depth, nor between R^2^ and genome coverage > 20X (Spearman’s, *p* > 0.05).

There was no significant difference in RMSE or R^2^ by template concentration (Kruskal-Wallis, *p* > 0.05, Supplemental Table 11-13). Differences in the R^2^ were noted, however, between mock communities of different compositions (Kruskal-Wallis, *p* = 0.03, Supplemental Table 11-13), specific differences could not be determined using post-hoc analyses (Dunn’s pairwise test, *p_H_*_olm_ > 0.05, Supplemental Table 11-13). The RMSE values were not significantly different between the different mock communities (Kruskal-Wallis, *p* = 0.26).

### Subset Data

Filtered reads were subset from each library that was sequenced with PE150 reads and PE300 reads to determine the threshold where sequencing depth is insufficient to correctly assign variants and estimate abundance. From the subset data, the resulting R^2^ values ranged from -1.43 to 1.00, and the RMSE values ranged from 0.64 to 22.0. Based on the global Friedman test, the R^2^ values were significantly affected by sequencing depth (Friedman’s tests, *p* < 0.01). Post-hoc analyses, however, only indicated a significant effect of sequencing depth for full-length VarSkip libraries sequenced with PE300 reads. Libraries subset to the lowest coverage (1X) were significantly less accurate than libraries subset to the highest coverages (10X, 12X, 25X, 50X, 100X, Dunn’s pairwise tests, *p_Holm_* < 0.01, Figure 3).

**Figure 3.**
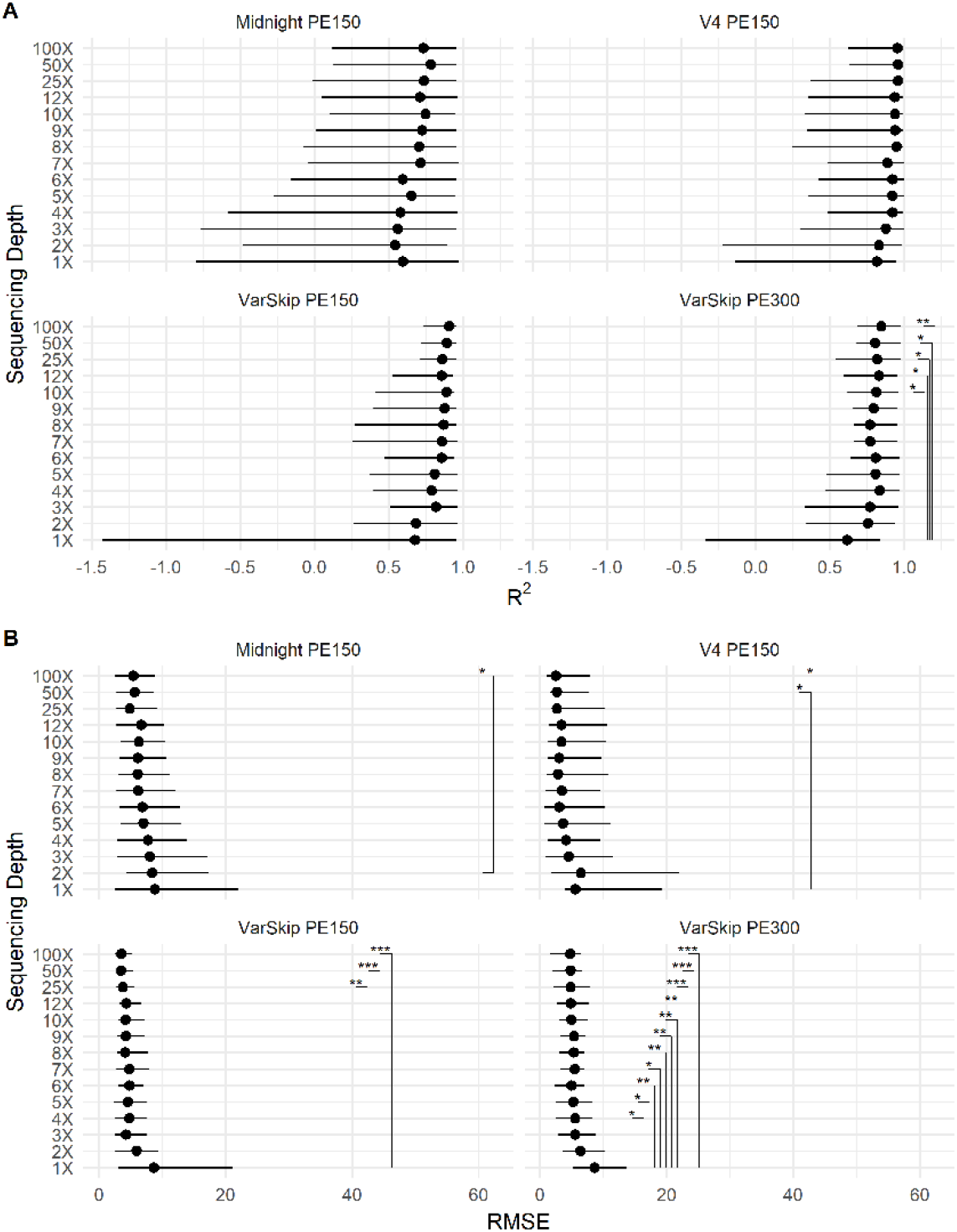
The effect of subsetting on the accuracy of estimates of variant abundance. Filtered reads were subset from libraries sequenced with PE150 reads and PE300, across a range of expected sequencing depths. Subset reads were analyzed by kallisto to estimate variant abundance. The accuracy of these estimates was assessed as the R^2^ value of the one-to-one model between expected abundance and observed abundance of the SARS-CoV-2 variants in each sample, and as the root-mean-squared error (RMSE) between expected and observed abundance. The median R^2^ and RMSE values are presented for each subset, along with the observed range of values.

The RMSE values were significantly affected by the depth of coverage (Friedman’s tests, *p* < 0.001). The Midnight libraries subset to 2X were less accurate than libraries subset to the highest coverage, 100X (Dunn’s pairwise test, *p_Holm_ =* 0.05, Figure 3). The V4 libraries subset to 1X were less accurate than libraries subset to 50X and 100X (Dunn’s pairwise tests, *p_Holm_ <* 0.05, Figure 3). The VarSkip libraries, sequenced with PE150 reads, subset to 1X were less accurate than libraries subset to 25X, 50X, and 100X (Dunn’s pairwise tests, *p_Holm_ <* 0.05, Figure 3). The full-length VarSkip libraries, sequenced with PE300 reads, subset to 1X were less accurate than all libraries subset to 4X or greater (Dunn’s pairwise tests, *p_Holm_ <* 0.05, Figure 3).

## DISCUSSION

Monitoring SARS-CoV-2 variants in wastewater is a promising new tool for new variant identification and outbreak tracking, however, it is notoriously challenging to implement due to the overwhelming abundance of non-target nucleic acids present in the complex wastewater matrix. For wastewater surveillance to be successful, sequencing of SARS-CoV-2 RNA requires genomic enrichment. Our results demonstrate that multiplexed tiled amplicon enrichment of SARS-CoV-2 in wastewater is a promising strategy for surveillance of SARS-CoV-2. We found that ARTIC V4 and sheared NEB VarSkip workflows provided sufficient data for use with kallisto to call variants and estimate their abundance with a high degree of accuracy. With further optimization, wastewater sequencing with either ARTIC V4 or NEB VarSkip workflows are likely to provide robust information for the genomic surveillance of variant(s) circulating within the population.

### Sequencing Coverage and Sequencing Depth

Obtaining high genome coverage is important for accurate variant identification and abundance estimations (Baaijens et al., 2021). SARS-CoV- 2 viral RNA is often found highly degraded and in low titers, requiring genomic enrichment prior to sequencing (Lin et al., 2021; Wurtzer et al., 2021). Even after amplicon enrichment, sequences obtained from wastewater often result in uneven coverage and depth across the SARS-CoV-2 genome (Lin et al., 2021; Smyth et al., 2022). We determined that libraries created with the NEB VarSkip library prep methods (both fragmented and full length) produced the highest genomic coverage and sequencing depth, when compared to ARTIC V4 and Midnight (Table 1). The starting concentrations of the mock community did not significantly impact the genomic coverage and sequencing depth acquired from these libraries.

Uneven genomic coverage was noted for all three tiled amplicon approaches but was especially apparent from libraries prepared with the 1,200-bp Freed/Midnight primer scheme. Regardless of sequencing effort, the Freed/Midnight consistently resulted in poor coverage and depth (Supplemental Figure 4). Several regions of the SARS-CoV-2 genome were entirely uncaptured by the Freed/Midnight amplicon tiled method. These gaps in coverage were likely due to the segmentation of the synthetic RNA controls, each of which are comprised of six 5-kp non-overlapping fragments (Twist Biosciences, 2022). Given this context, these data demonstrate that long amplicons are not appropriate for sequencing genomic material that is fragmented and heavily degraded, as we would expect of SARS-CoV-2 RNA isolated from wastewater. Our findings are consistent with those by Lin et al. (2021), who found that shorter amplicons are more resilient to sample RNA degradation than the larger Freed/Midnight amplicons.

### SARS-CoV-2 Variant Identification from Wastewater

Variant identification was performed via a pipeline containing kallisto as described by Baaijens, et al. (2021). With this computational approach, target SARS-CoV-2 variants were called from all libraries containing mock communities (Supplemental Figure 7). Even variants that were at low proportions within the mock community (3% and 14%), were detected in all libraries. These results demonstrate better variant calling than Baaijens et al. (2021), who found it challenging to detect variants in wastewater with clinical frequencies less than 10%. This suggests high-quality amplicon libraries can detect the emergence of low-frequency variants, overcoming a significant limitation for use in public health surveillance.

While the sheared VarSkip libraries contained the most sequencing reads from the wastewater mock communities assigned to reference genomes by kallisto, the ARTIC V4 libraries had the most accurate assignments of the mock community reads (Figure 2). Often, mock community reads were assigned to off-target variants that were not spiked into the original sample but were present in the reference database. These off-target calls result in noise that can obscurethe true dynamics of circulating variants. To reduce noise, Baaijens et al. (2021) recommended filtering results by applying a minimal abundance threshold of 0.1%. Even when filtering for low-abundant calls, we detected off-target variants ubiquitously across the samples. However, in the ARTIC V4 libraries we saw low proportions of off target reads being called (<2%) and accurate variant calls for the reads that were assigned to genomes in the reference database (Figure 2). This was observed with both read lengths and across all mock community starting concentrations.

### Predictions of Variant Abundance

The sequencing and analysis of wastewater mock communities resulted in highly accurate abundance estimates of SARS-CoV-2 variants, with R^2^ values reaching up to 0.99 (Supplemental Figure 11-13). Libraries prepared with ARTIC V4 and NEB VarSkip workflows resulted in significantly better estimates than libraries prepared with the Freed/Midnight amplicons, with ARTIC V4 libraries making the most accurate estimations out of all the library preparation methods (Figure 2A). This accuracy seen in the estimations made from ARTIC V4 libraries can be seen in both sequencing read lengths. These results suggest that library preparation with the ARTIC V4 workflow may be the most efficient approach to genomic surveillance for SARS-CoV-2 variants from wastewater. The NEB VarSkip and Freed/Midnight approaches require fragmentation if they are to be run on an Illumina machine that yields PE150 or PE250 reads, whereas ARTIC V4 amplicons do not require end preparation with fragmentation. Additionally, ARTIC V4 amplicons can be sequenced with a read length of PE150, which are generally less expensive, without needing fragmentation. Therefore, we conclude that library preparation with the ARTIC V4 workflow is less expensive and more efficient than NEB VarSkip and Freed/Midnight library preparation methods.

### Threshold for Sequencing

Filtered reads were subset from each mock community library that was sequenced with PE150 reads and PE300 reads to determine the minimum sequencing requirements for obtaining accurate variant assignment and abundance estimations for each library type. There has been no obvious comparison of the minimum sequencing requirements needed from each of the library preparation methods described here. We determined that when using ARTIC V4 sequencing reads, there is no significant difference in the accuracy of variant detection and estimation between sequencing depths (from 1X to 100X) (Figures 3). This illustrates how low the sequencing depth of a sample prepared with ARTIC V4 libraries can be and still obtain the correct variant identifications and estimations. Similarly, subset reads from fragmented VarSkip libraries had no significant differences between any of the sequencing depths when using the R^2^ to measure accurate identification and estimation. However, when using the RMSE values there were significant differences seen between 1X of sequencing depth and a few of the higher sequencing depths (Figures 3). From this we determine that fragmented VarSkip libraries can also have moderately low sequencing depth (≥2X) and still obtain accurate results. The same cannot be said for libraries prepared with the Freed/Midnight amplicons or libraries prepared with full length VarSkip; both of which had higher R^2^ for all coverage depths and significant differences between many of the coverage depths (Figure 3).

## CONCLUSION

Several sequencing methods have been proposed for genomic surveillance of SARS-CoV-2 in wastewater, but there is little consensus on the most appropriate approaches for variant identification and abundance estimations. By generating, sequencing, and evaluating a series of mock wastewater communities, we directly compared three tiled amplicon approaches for the whole genome enrichment of SARS-CoV-2 variants from wastewater. We demonstrate that while mock community reads created with NEB VarSkip library preparation methods yield the highest genomic coverage and sequencing depth, ARTIC V4 libraries obtain the most accurate variant identifications and abundance estimations. We also illustrate that mock community reads created with the Freed/Midnight library preparation method are not ideal for variant detection or abundance estimation when using short read sequencing. In summary, ARTIC V4 and NEB VarSkip approaches are applicable for the routine monitoring of circulating SARS-CoV-2 variants from wastewater. To perform this task in the most labor- and cost-efficient way, we recommend using the ARTIC V4 library preparation methods.

## METHODS

### Background Wastewater Matrix

Time-composite (24 h) wastewater influent samples were collected from Athens-Clarke County, Georgia (USA) over multiple days and three separate wastewater treatment plants in May 2020, when viral titers of SARS-CoV-2 were below the limit of detection (Lott et al., 2023). Bulk wastewater samples were stored at –20°C prior to total nucleic acid extraction. DNA and RNA were extracted from wastewater samples were extracted using the Zymo Environ Water RNA Kit (Cat No. R2042, Zymo Research Corp, Irvine, CA). Briefly, DNA and RNA were extracted from 10 mL aliquots of wastewater according to the manufacturer’s instructions. Extracted RNA and DNA were eluted in 60 μL of molecular-grade water. 84 aliquots of influent wastewater were processed, and the eluates were combined for a total volume of 5 mL of DNA/RNA extract, which served as the diluent (background) in the preparation of the mock communities. The background matrix was stored at –20°C prior to preparation with mock communities.

### Mock Communities

Mock communities were prepared by spiking synthetic genomic RNA of known SARS-CoV-2 variants into the background wastewater matrix. Synthetic genomic viral RNA for each of four SARS-CoV-2 variants - Wuhan, Alpha (B.1.1.7), Beta (B.1.351), and Delta (AY.2) - were sourced from Twist Biosciences (Supplemental Table 2) and combined at varying proportions to prepare five distinct mock communities (Table 2). Mock communities were diluted into the background wastewater matrix to a final concentration of 10^3^, 10^2^, and 10^1^ copies μL (all variants combined).

**Table 2.** Wastewater Mock Community SARS-CoV-2 Variant Composition. Synthetic Genomic RNA for SARS-CoV-2 variants were sourced from Twist Biosciences and combined at known proportions to prepare five distinct mock community profiles. Mock communities were then diluted into the background wastewater matrix to a final concentration of 10^3^, 10^2^, and 10^1^ copies μL^-1^, resulting in fifteen unique mock communities.

| Mock Community No. | Wuhan | Alpha (B.1.1.7) | Beta (B.1351) | Delta (AY.2) |
| --- | --- | --- | --- | --- |
| **1** | 3% | 14% | 28% | 55% |
| **2** | 55% | 3% | 14% | 28% |
| **3** | 28% | 55% | 3% | 14% |
| **4** | 14% | 28% | 55% | 3% |
| **5** | 25% | 25% | 25% | 25% |

### Library Preparation

Immediately following mock community preparation, cDNA was synthesized from each sample using LunaScript® RT SuperMix (Cat No. E3010, New England BioLabs, Ipswich, MA), according to the manufacturer’s recommended reaction conditions (Supplemental Table 3). The cDNA was stored at –20°C prior to whole genome enrichment with either ARTIC V4, VarSkip, or Midnight primer sets.

Midnight amplicon libraries were prepared with the Freed/Midnight Amplicon Panel workflow, modified for sequencing with Illumina chemistry. Briefly, 1,200 bp amplicons were generated with the Freed/Midnight primer pools (Freed, Vlková, Faisal, & Silander, 2021, Cat No. 10011644) and Q5® Hot Start High-Fidelity 2X Master Mix, according to recommended reaction conditions (Supplemental Table 4). The two amplicon pools were combined in equal proportions and bead cleaned at 0.65X. Fragmentation and end repair were conducted with NEBNext® End Prep Ultra II FS reagents (Supplemental Table 4, Cat No. E7805, Ipswich, MA). Custom iTru adapters were ligated to amplicons with NEBNext® Ultra II Ligation Master Mix (Supplemental Table 6). A second bead clean-up was performed at 0.8X before proceeding with a final PCR enrichment with iTru indices (Supplemental Table 7).

NEB VarSkip amplicon libraries were prepared according to the NEBNext® ARTIC SARS-CoV-2 FS Library Prep Kit Workflow for Illumina® (NEB, Cat No. E7658, Ipswich, MA).

Briefly, 500 bp amplicons were generated using NEB VarSkip Short primer pools (Gautreau, 2021, NEB, Cat No. E7658) and Q5® HotStart MasterMix, under recommended reaction conditions (Supplemental Table 8). The two amplicon pools were combined in equal proportions and bead cleaned at 0.8X. Amplicons prepared for short-read Illumina sequencing (PE150 and PE250 reads) were fragmented with NEBNext® End Prep Ultra II FS reagents (Supplemental Table 5, Cat No. E7805, Ipswich, MA). Amplicons prepared for long-Illumina read sequencing (PE300) underwent end repair with NEBNext® End Prep Ultra II Kit (Supplemental Table 9, Cat No. E7645, NEB, Ipswich, MA). Custom iTru adapters were ligated to amplicons with NEBNext® Ultra II Ligation Master Mix with a 15 min incubation at 20°C (Supplemental Table 6). A second bead clean-up was performed at 0.8X before proceeding with PCR enrichment with iTru indices (Supplemental Table 7).

ARTIC V4 amplicon libraries were prepared according to the NEBNext® ARTIC SARS-CoV-2 Library Prep Kit Workflow for Illumina® (Discontinued, New England BioLabs, Ipswich, MA). Briefly, 400 bp amplicons were generated using ARTIC V.4.0 primer pools (Quick, 2020), and Q5® Hot Start High-Fidelity 2X Master Mix (NEB, Cat No. M0494, Ipswich, MA), according to the recommended reaction conditions (Supplemental 8). The two amplicon pools were combined in equal proportions and cleaned with a 0.8X SpeedBead™ clean-up (Sera-Mag SpeedBeads™, Cat No. 65152105050250, Cytiva, USA). Following end repair with NEBNext® End Prep Ultra II Kit (Supplemental Table 9, Cat No. E7645, NEB, Ipswich, MA), custom iTru adapters were ligated to amplicons with NEBNext® Ultra II Ligation Master Mix (Cat No. E7648, NEB, Ipswich, MA) with a 15 min incubation at 20°C (Supplemental Table 8). A second clean-up was performed at 0.9X before proceeding with PCR enrichment with iTru indices (Supplemental Table 7).

All adapter-ligated amplicons were amplified with custom iTru5 and iTru7 index primers (Glenn *et al*. 2019) with NEBNext Ultra II Q5 Master Mix (NEB, Cat No, M0544, Ipswich, MA) according to reaction conditions described in Supplemental Table 6. A final bead clean was used to select NEB VarSkip full length amplicons ∼500 bp, ARTIC V4 amplicons ∼400 bp, NEB VarSkip sheared amplicons ∼300 bp, and Midnight amplicons ∼ 300bp for sequencing on Illumina MiSeq with PE250 reads (500 cycles) and on Illumina HiSeq with PE150 reads (300 cycles). Unfragmented NEB VarSkip amplicons ∼500 bp were sequenced on Illumina MiSeq with PE300 reads (600 cycles). Libraries were sequenced to target 10^5^ PE150 reads, 10^3^ PE250 reads, and 10^4^ PE300 reads.

### Controls

Several positive and negative controls were prepared as amplicon libraries and sequenced in parallel with mock communities. Positive extraction controls included a heat-inactivated strain of the SARS-CoV-2 Wuhan variant carried in 1X phosphate buffered saline (PBS) and a heat-inactivated strain of the SARS-CoV-2 Wuhan variant, carried in wastewater (ATCC VR-1986HK). Viral RNA was extracted from these positive controls using the Zymo Environ Water Kit, as described previously, and prepared as amplicon libraries at a starting concentration of approximately 150 copies μL . Alongside these extraction controls, the Twist Wuhan control was prepared into amplicon libraries at a starting concentration of 10^3^ copies L^-1^. Negative controls included the background wastewater matrix in addition to molecular-grade water.

### Sequence Cleaning

Trimmomatic v0.39 was used to perform quality filtering and to remove adapters from the raw reads (Bolger, Lohse, & Usadel, 2014). Leading and trailing parameters were set to quality of three Phred-33 to remove low quality bases from the reads. The sliding window parameter was set to 4:20 to quality trim the reads and the minlen parameter was set to exclude sequences that were less than 100 bp. To quantify coverage, sequences were aligned to the SARS-CoV-2 Wuhan reference genome (GenBank MN908947) using bbmap (Bushnell, 2014).

### Variant Abundance Estimation

Kallisto was used to assign variants and estimate their abundance within a sample (Bray, Pimentel, Melsted, & Pachter, 2016). The reference database for kallisto was created using high-quality genomes obtained from GISAID (Elbe & BucklanduMerrett, 2017). Genomes that were categorized as high quality, high coverage, no gaps and low number of N’s were used in the reference database. Whenever possible, five genomes were used for each known variant of concern or variant of interest within the population. A list of variants and the reference genomes used for each variant can be found in Supplemental Table 10. Trimmed and cleaned reads were used to estimate the abundance of SARS-CoV-2 variants which were estimated using the kallisto quant command with the –b flag set to 100 (Baaijens et al., 2021). Variant abundance estimations were not adjusted for noise seen in the negative controls, though a minimum abundance threshold of 0.1% was applied to minimize false positives.

### Statistical Analyses and Data Visualization

Sequencing statistics, genome coverage, and SARS-CoV-2 variant abundance estimates are provided in Supplementary Data. Raw data were examined and visualized using R v.4.3.1 in RStudio v.2022.07.2+576. Analysis scripts are available on Github (https://github.com/meganejlott/ww_mock_community). Additional Details for Statistical Analyses are described in Supplemental Text.

### Subsetting Filtered Sequence Data

The trimmed reads for all samples with HiSeq PE150 read and MiSeq PE300 reads were subset using seqtk v1.4 (Li, 2012). For all mock community libraries, the trimmed reads were subset to create coverages of 1X, 2X, 3X, 4X, 5X, 6X, 7X, 8X, 9X, 10X, 12X, 25X, 50X, and 100X. After subsetting, the subset samples for each sample type went through the same variant calling process the original samples were processed through.

## AUTHORSHIP

Conceptualization: TCG, EKL; Methodology: MEJL, AHS, LML, MSB; Investigation: MEJL, AHS, LML, KCD; Formal analysis: AHS, MEJL, WAN; Writing - review and editing: AHS, MEJL, LML, WAN, KCD, MSB, TCG, EKL; Funding acquisition: EKL; Supervision: EKL, TCG.

## ACKNOWLEDGEMENTS

Funding in support of this work was provided by U.S. Centers for Disease Control and Prevention through Contract No. 75D30121C11163 to EKL.

