## Supplemental Tables and Figures for "Comparison of three tiled amplicon sequencing approaches for SARS-CoV-2 variant detection from wastewater"

**SUPPLEMENTARY TABLES AND FIGURES**

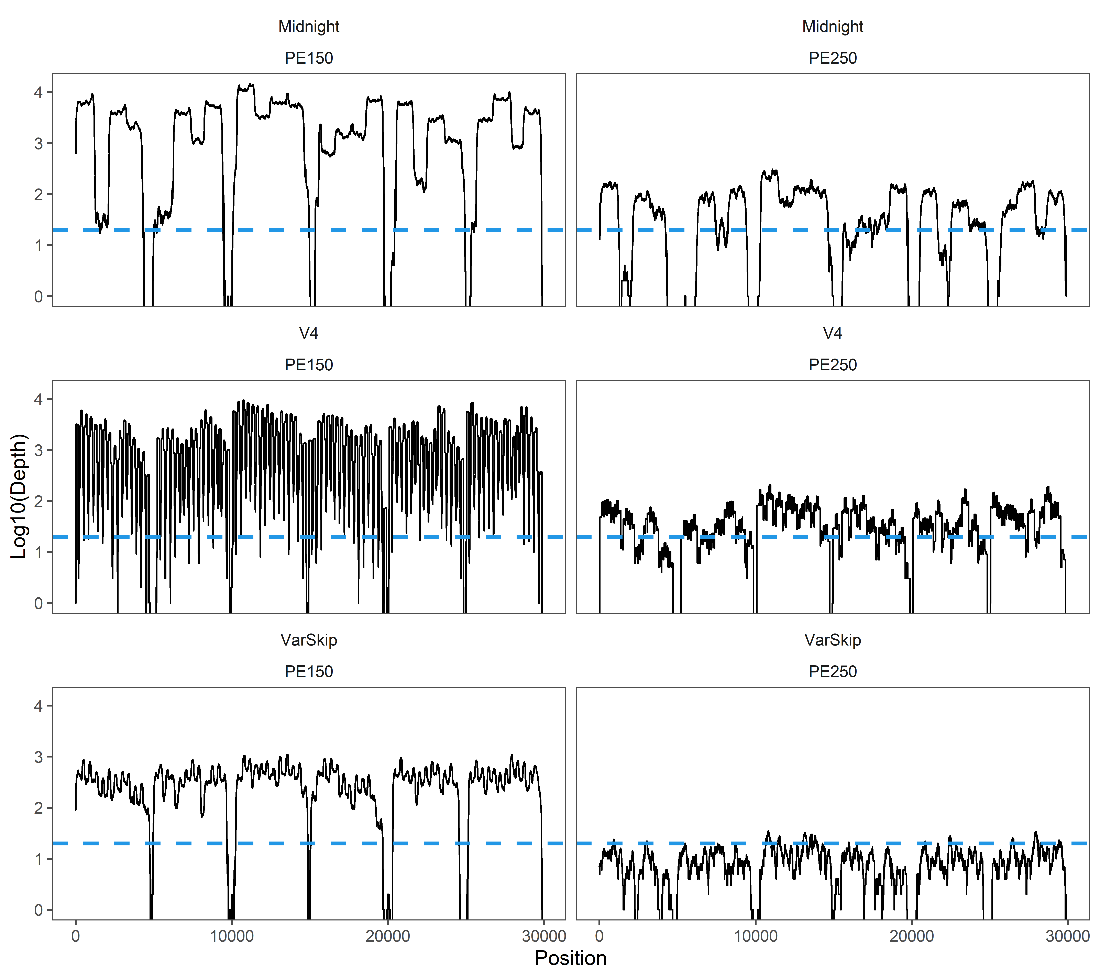

**Supplemental Figure 1. Genomic coverage of the SARS-CoV-2 Wuhan Twist Control using three tiled amplicon preparation methods.** The SARS-CoV-2 Wuhan Twist Control was enriched using the Freed/Midnight, ARTIC V4, and NEB VarSkip workflows. Libraries were sequenced with Illumina PE150 and PE250 reads, with approximately 10^3^ and 10^6^ reads, respectively. The resulting depth figures are annotated with a dashed line representing the threshold for 20X coverage.

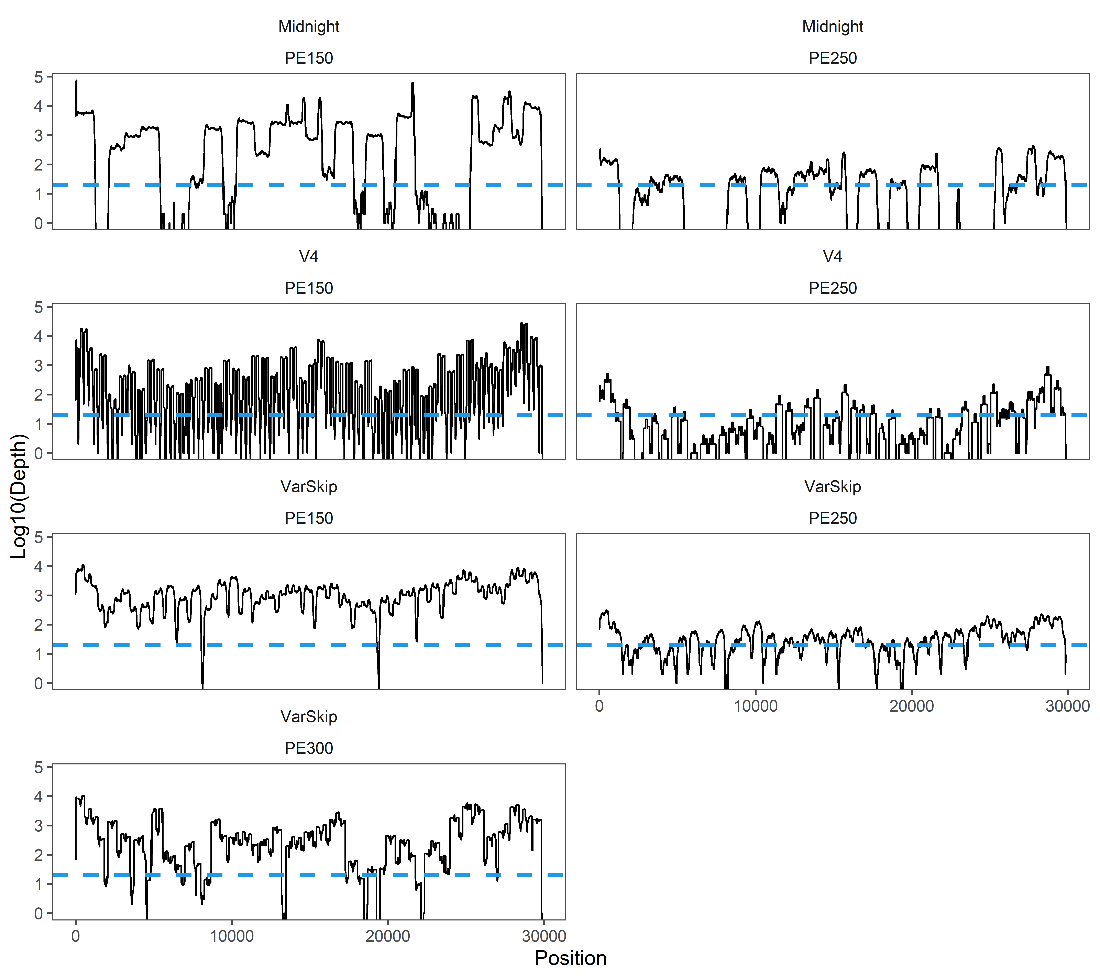

**Supplemental Figure 2.** **Genomic coverage of the heat-inactivated SARS-CoV-2 control, carried in PBS, using three tiled amplicon preparation methods.** The heat-inactivated SARS-CoV-2 control, carried in PBS, was extracted using the Zymo Environmental RNA Kit. Extracted genomic material was then enriched using the Freed/Midnight, ARTIC V4, and NEB VarSkip workflows. Libraries were sequenced with Illumina PE150, PE250 and PE300 reads, with approximately 10^3^ and 10^6^ reads, respectively. The resulting depth figures are annotated with a dashed line representing the threshold for 20X coverage.

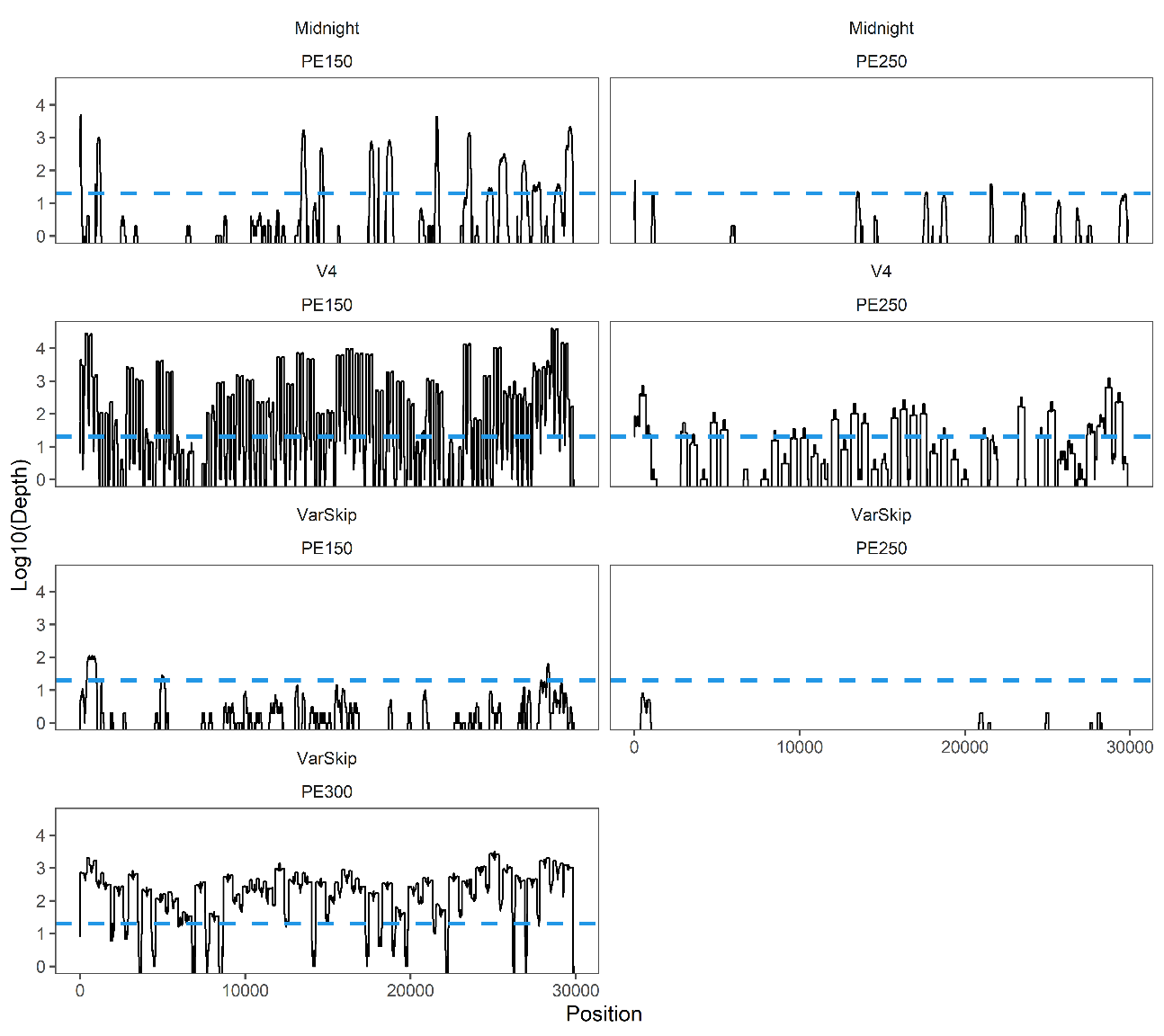

**Supplemental Figure 3. Genomic coverage of the heat-inactivated SARS-CoV-2 control, carried in wastewater, using three tiled amplicon preparation methods.** The heat-inactivated SARS-CoV-2 control, carried in wastewater, was extracted using the Zymo Environmental RNA Kit. Extracted genomic material was then enriched using the Freed/Midnight, ARTIC V4, and NEB VarSkip workflows. Libraries were sequenced with Illumina PE150 and PE250 reads, with approximately 10^3^ and 10^5^ reads, respectively. The resulting depth figures are annotated with a dashed line representing the threshold for 20X coverage.

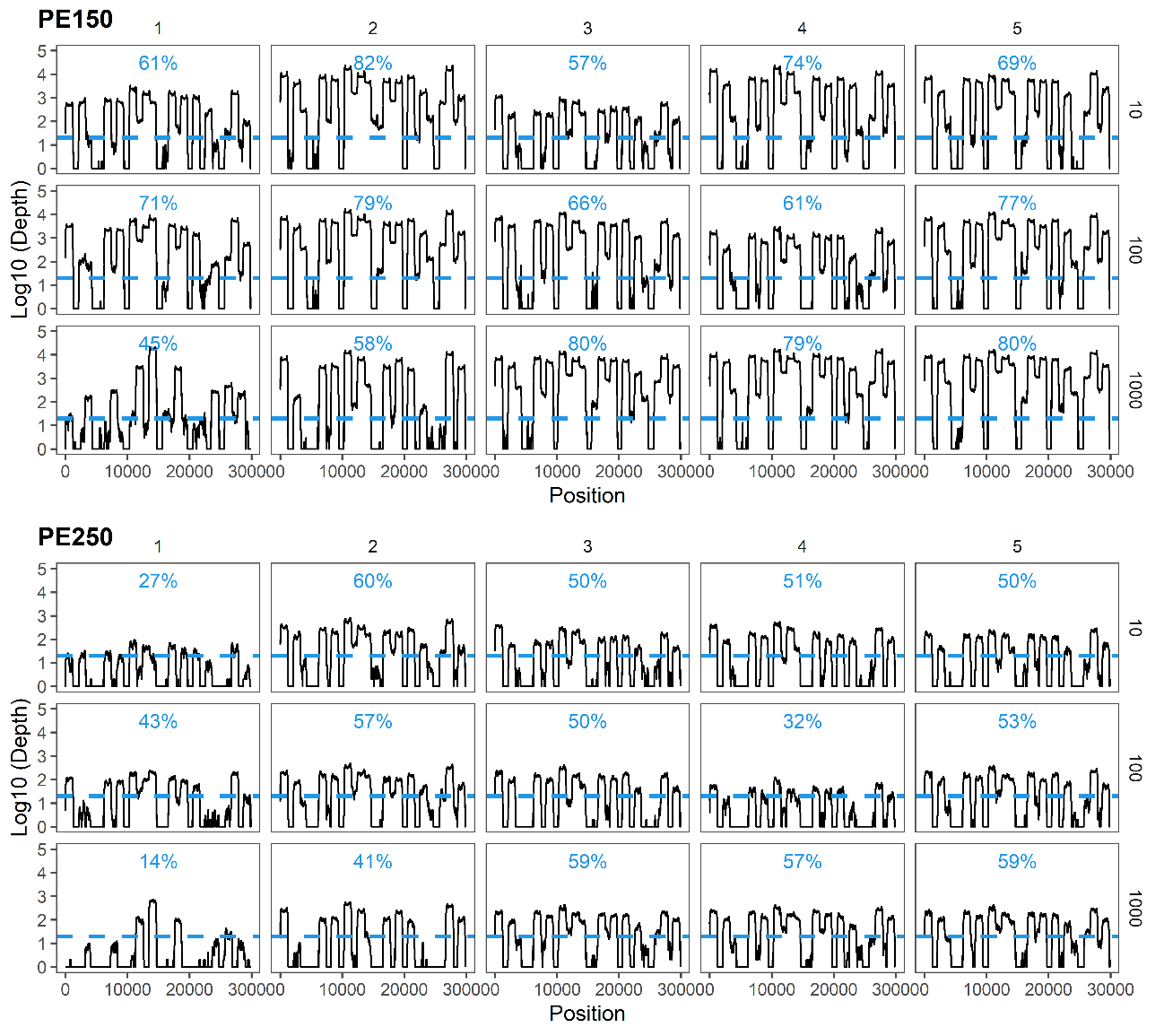

**Supplemental Figure 4. Genomic coverage of the wastewater mock communities, using the Freed/Midnight sequencing workflow.** Synthetic genomic RNA for SARS-CoV-2 variants were combined at varying proportions to prepare five district mock communities (numbered 1 through 5), and diluted into extracted wastewater to concentrations of 10^3^, 10^2^, and 10^1^ copies μL^-1^. Mock communities were then enriched using the Freed/Midnight workflow and libraries were sequenced with Illumina PE150 and PE250 chemistry, with approximately 10^3^ and 10^5^ reads, respectively. The resulting depth figures are marked with a dashed line representing the threshold for 20X coverage and annotated with the percentage of the genome sequenced at or beyond this threshold.

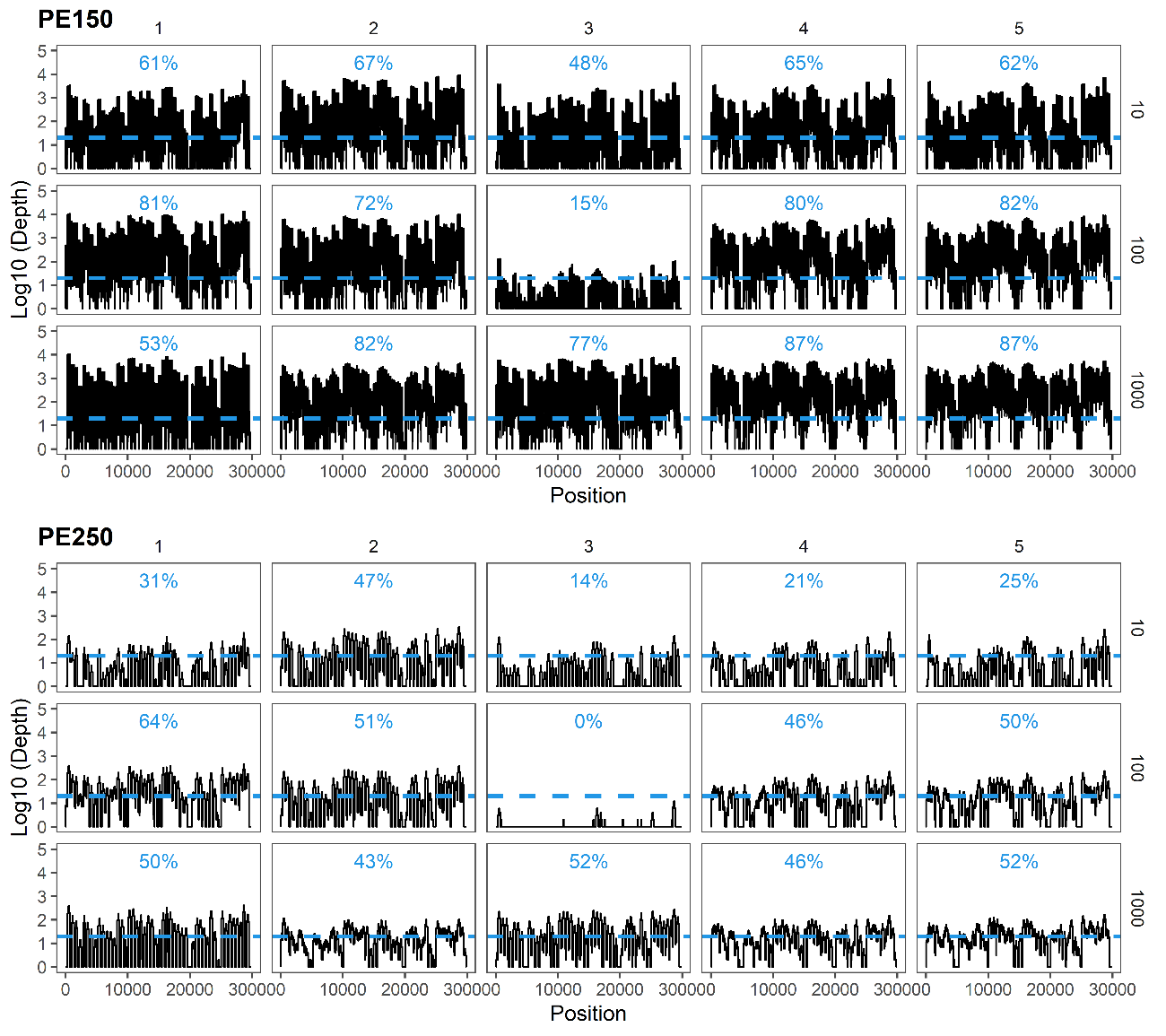

**Supplemental Figure 5. Genomic coverage of the wastewater mock communities, using the ARTIC V4 sequencing workflow.** Synthetic genomic RNA for SARS-CoV-2 variants were combined at varying proportions to prepare five district mock communities (numbered 1 through 5), and diluted into extracted wastewater to concentrations of 10^3^, 10^2^, and 10^1^ copies μL^-1^. Mock communities were then enriched using the ARTIC V4 workflow and libraries were sequenced with Illumina PE150 and PE250 chemistry, with approximately 10^3^ and 10^5^ reads, respectively. The resulting depth figures are marked with a dashed line representing the threshold for 20X coverage and annotated with the percentage of the genome sequenced at or beyond this threshold.

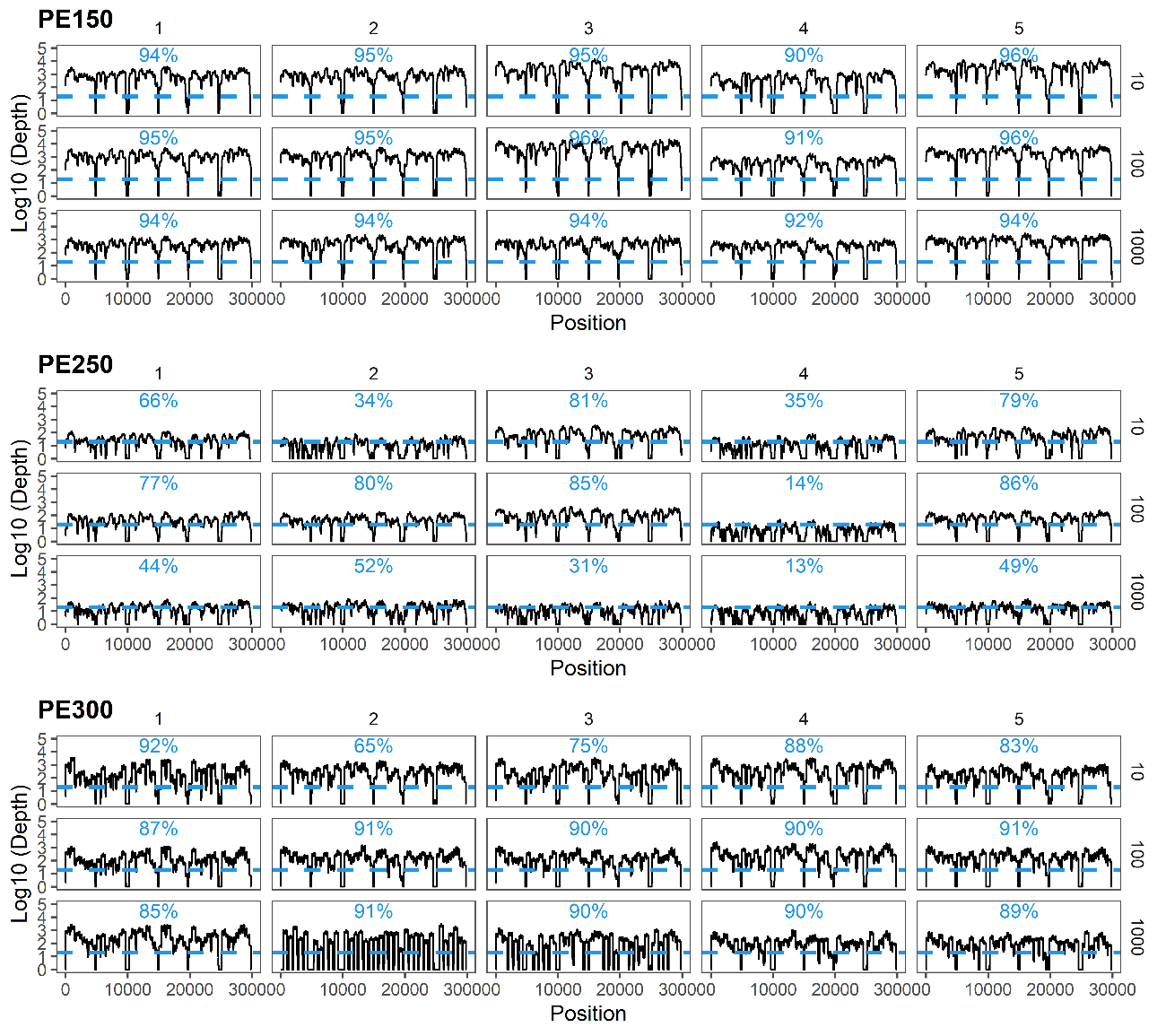

**Supplemental Figure 6. Genomic coverage of the wastewater mock communities, using the NEB VarSkip sequencing workflow.** Synthetic genomic RNA for SARS-CoV-2 variants were combined at varying proportions to prepare five district mock communities (numbered 1 through 5), and diluted into extracted wastewater to concentrations of 10^3^, 10^2^, and 10^1^ copies μL^-1^. Mock communities were then enriched using the NEB VarSkip workflow and libraries were sequenced with Illumina PE150 and PE250 chemistry, with approximately 10^3^ and 10^5^ reads, respectively. The resulting depth figures are marked with a dashed line representing the threshold for 20X coverage and annotated with the percentage of the genome sequenced at or beyond this threshold.

**Supplemental Table 1. Target and non-target detection of SARS-CoV-2 variants, as assigned by kallisto.** Wastewater mock communities prepared with four known variants of SARS-CoV-2, the parent Wuhan lineage, as well as the Alpha, Beta, Delta lineages, spiked at known proportions. Sequencing reads were assessed using the kallisto workflow to assign reads to SARS-CoV-2 variants in the reference set. The reference set included genomes from four the target variants as well as genomes from eleven off-target variants.

| Sequencing Approach | | WHO Name | Detection Observed, %  (No. Samples Detected/  No. Samples Total) | Detection Expected, %  (No. Samples Detected/  No. Samples Total) |
| --- | --- | --- | --- | --- |
| Midnight PE150 | | Wuhan | 100% (15/15) | 100% (15/15) |
|  | Alpha | | 100% (15/15) | 100% (15/15) |
|  | Beta | | 100% (15/15) | 100% (15/15) |
|  | Delta | | 100% (15/15) | 100% (15/15) |
|  | Epsilon | | 100% (15/15) | 0% (0/15) |
|  | Eta | | 93% (14/15) | 0% (0/15) |
|  | Gamma | | 93% (14/15) | 0% (0/15) |
|  | Iota | | 100% (15/15) | 0% (0/15) |
|  | Kappa | | 100% (15/15) | 0% (0/15) |
|  | Mu | | 100% (15/15) | 0% (0/15) |
|  | Omicron_BA.1 | | 100% (15/15) | 0% (0/15) |
|  | Omicron_BA.2 | | 93% (14/15) | 0% (0/15) |
|  | Omicron_BA.4 | | 87% (13/15) | 0% (0/15) |
|  | Omicron_BA.5 | | 100% (15/15) | 0% (0/15) |
|  | Zeta | | 73% (11/15) | 0% (0/15) |
| Midnight PE250 | | Wuhan | 100% (15/15) | 100% (15/15) |
|  | Alpha | | 100% (15/15) | 100% (15/15) |
|  | Beta | | 100% (15/15) | 100% (15/15) |
|  | Delta | | 100% (15/15) | 100% (15/15) |
|  | Epsilon | | 100% (15/15) | 0% (0/15) |
|  | Eta | | 73% (11/15) | 0% (0/15) |
|  | Gamma | | 100% (15/15) | 0% (0/15) |
|  | Iota | | 100% (15/15) | 0% (0/15) |
|  | Kappa | | 93% (14/15) | 0% (0/15) |
|  | Mu | | 93% (14/15) | 0% (0/15) |
|  | Omicron_BA.1 | | 100% (15/15) | 0% (0/15) |
|  | Omicron_BA.2 | | 100% (15/15) | 0% (0/15) |
|  | Omicron_BA.4 | | 87% (13/15) | 0% (0/15) |
|  | Omicron_BA.5 | | 93% (14/15) | 0% (0/15) |
|  | Zeta | | 67% (10/15) | 0% (0/15) |
| V4 PE150 | | Wuhan | 100% (15/15) | 100% (15/15) |
|  | Alpha | | 100% (15/15) | 100% (15/15) |
|  | Beta | | 100% (15/15) | 100% (15/15) |
|  | Delta | | 100% (15/15) | 100% (15/15) |
|  | Epsilon | | 20% (3/15) | 0% (0/15) |
|  | Eta | | 7% (1/15) | 0% (0/15) |
|  | Gamma | | 0% (0/15) | 0% (0/15) |
|  | Iota | | 7% (1/15) | 0% (0/15) |
|  | Kappa | | 93% (14/15) | 0% (0/15) |
|  | Mu | | 100% (15/15) | 0% (0/15) |
|  | Omicron_BA.1 | | 13% (2/15) | 0% (0/15) |
|  | Omicron_BA.2 | | 7% (1/15) | 0% (0/15) |
|  | Omicron_BA.4 | | 0% (0/15) | 0% (0/15) |
|  | Omicron_BA.5 | | 0% (0/15) | 0% (0/15) |
|  | Zeta | | 0% (0/15) | 0% (0/15) |
| V4 PE250 | | Wuhan | 100% (15/15) | 100% (15/15) |
|  | Alpha | | 100% (15/15) | 100% (15/15) |
|  | Beta | | 100% (15/15) | 100% (15/15) |
|  | Delta | | 100% (15/15) | 100% (15/15) |
|  | Epsilon | | 60% (9/15) | 0% (0/15) |
|  | Eta | | 0% (0/15) | 0% (0/15) |
|  | Gamma | | 0% (0/15) | 0% (0/15) |
|  | Iota | | 87% (13/15) | 0% (0/15) |
|  | Kappa | | 87% (13/15) | 0% (0/15) |
|  | Mu | | 80% (12/15) | 0% (0/15) |
|  | Omicron_BA.1 | | 27% (4/15) | 0% (0/15) |
|  | Omicron_BA.2 | | 7% (1/15) | 0% (0/15) |
|  | Omicron_BA.4 | | 0% (0/15) | 0% (0/15) |
|  | Omicron_BA.5 | | 60% (9/15) | 0% (0/15) |
|  | Wuhan | | 100% (15/15) | 0% (0/15) |
|  | Zeta | | 0% (0/15) | 0% (0/15) |
| VarSkip PE150 | | Wuhan | 100% (15/15) | 100% (15/15) |
|  | Alpha | | 100% (15/15) | 100% (15/15) |
|  | Beta | | 100% (15/15) | 100% (15/15) |
|  | Delta | | 100% (15/15) | 100% (15/15) |
|  | Epsilon | | 100% (15/15) | 0% (0/15) |
|  | Eta | | 93% (14/15) | 0% (0/15) |
|  | Gamma | | 100% (15/15) | 0% (0/15) |
|  | Iota | | 100% (15/15) | 0% (0/15) |
|  | Kappa | | 100% (15/15) | 0% (0/15) |
|  | Mu | | 100% (15/15) | 0% (0/15) |
|  | Omicron_BA.1 | | 100% (15/15) | 0% (0/15) |
|  | Omicron_BA.2 | | 100% (15/15) | 0% (0/15) |
|  | Omicron_BA.4 | | 100% (15/15) | 0% (0/15) |
|  | Omicron_BA.5 | | 100% (15/15) | 0% (0/15) |
|  | Zeta | | 100% (15/15) | 0% (0/15) |
| VarSkip PE250 | | Wuhan | 100% (15/15) | 100% (15/15) |
|  | Alpha | | 100% (15/15) | 100% (15/15) |
|  | Beta | | 100% (15/15) | 100% (15/15) |
|  | Delta | | 100% (15/15) | 100% (15/15) |
|  | Epsilon | | 100% (15/15) | 0% (0/15) |
|  | Eta | | 60% (9/15) | 0% (0/15) |
|  | Gamma | | 100% (15/15) | 0% (0/15) |
|  | Iota | | 100% (15/15) | 0% (0/15) |
|  | Kappa | | 100% (15/15) | 0% (0/15) |
|  | Mu | | 100% (15/15) | 0% (0/15) |
|  | Omicron_BA.1 | | 100% (15/15) | 0% (0/15) |
|  | Omicron_BA.2 | | 93% (14/15) | 0% (0/15) |
|  | Omicron_BA.4 | | 100% (15/15) | 0% (0/15) |
|  | Omicron_BA.5 | | 100% (15/15) | 0% (0/15) |
|  | Zeta | | 100% (15/15) | 0% (0/15) |
| VarSkip PE300 | | Wuhan | 100% (15/15) | 100% (15/15) |
|  | Alpha | | 100% (15/15) | 100% (15/15) |
|  | Beta | | 100% (15/15) | 100% (15/15) |
|  | Delta | | 100% (15/15) | 100% (15/15) |
|  | Epsilon | | 40% (6/15) | 0% (0/15) |
|  | Eta | | 0% (0/15) | 0% (0/15) |
|  | Gamma | | 100% (15/15) | 0% (0/15) |
|  | Iota | | 53% (8/15) | 0% (0/15) |
|  | Kappa | | 27% (4/15) | 0% (0/15) |
|  | Mu | | 60% (9/15) | 0% (0/15) |
|  | Omicron_BA.1 | | 20% (3/15) | 0% (0/15) |
|  | Omicron_BA.2 | | 0% (0/15) | 0% (0/15) |
|  | Omicron_BA.4 | | 0% (0/15) | 0% (0/15) |
|  | Omicron_BA.5 | | 7% (1/15) | 0% (0/15) |
|  | Zeta | | 93% (14/15) | 0% (0/15) |

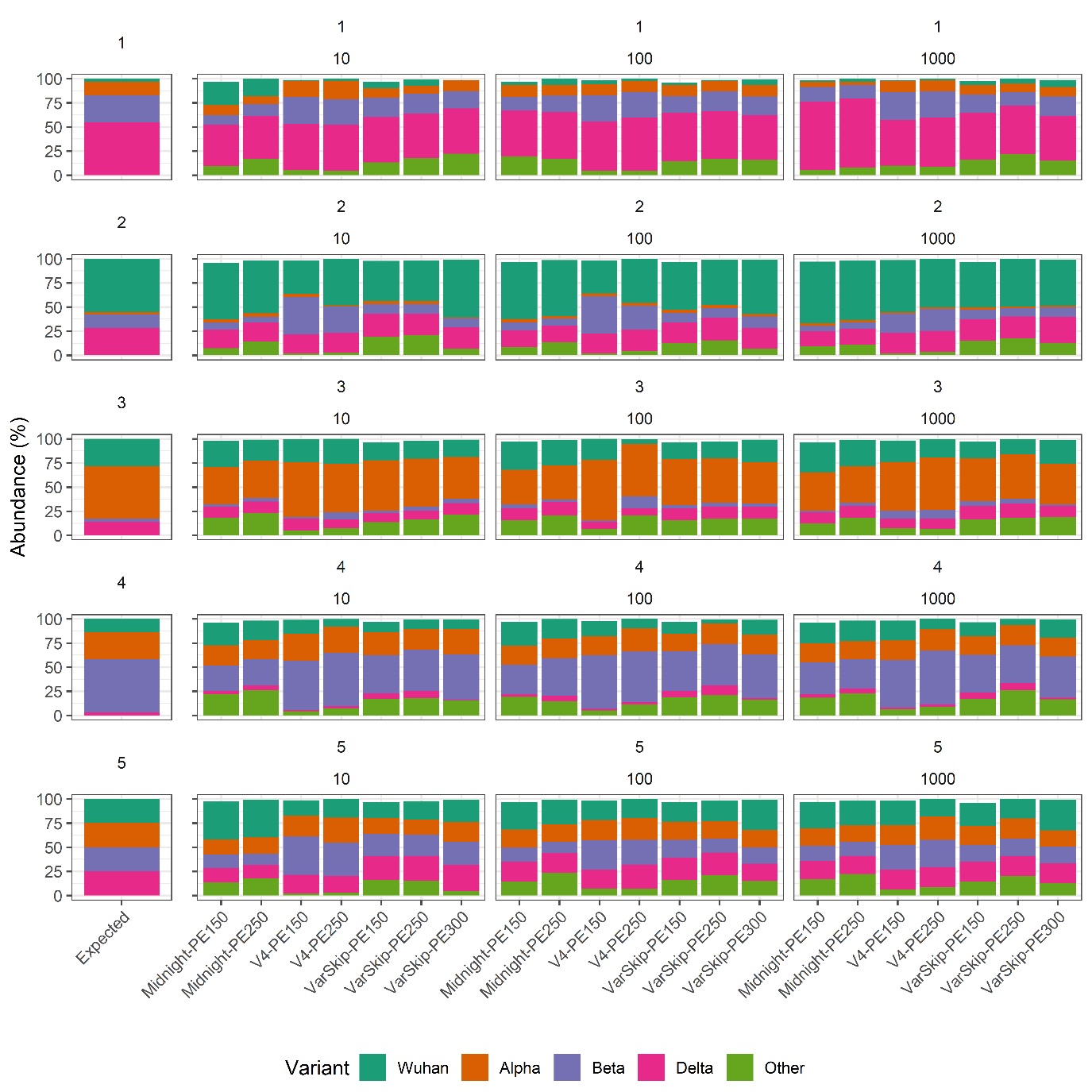

**Supplemental Figure 7.** **Abundance estimates from wastewater mock communities prepared with three whole-genome tiled amplicon enrichment methods.** Synthetic genomic RNA for SARS-CoV-2 variants were combined at varying proportions to prepare five district mock communities (numbered 1 through 5), and diluted into extracted wastewater to concentrations of 10^3^, 10^2^, and 10^1^ copies μL^-1^. Mock communities were then enriched as Freed/Midnight, ARTIC V4, and NEB VarSkip libraries and sequenced with Illumina PE250 and PE150 chemistry, with approximately 10^3^ and 10^5^ reads, respectively. Sequencing reads were assessed using the kallisto workflow to estimate abundance of SARS-CoV-2 variants. Expected abundance was compared to observed abundance, reported by kallisto.

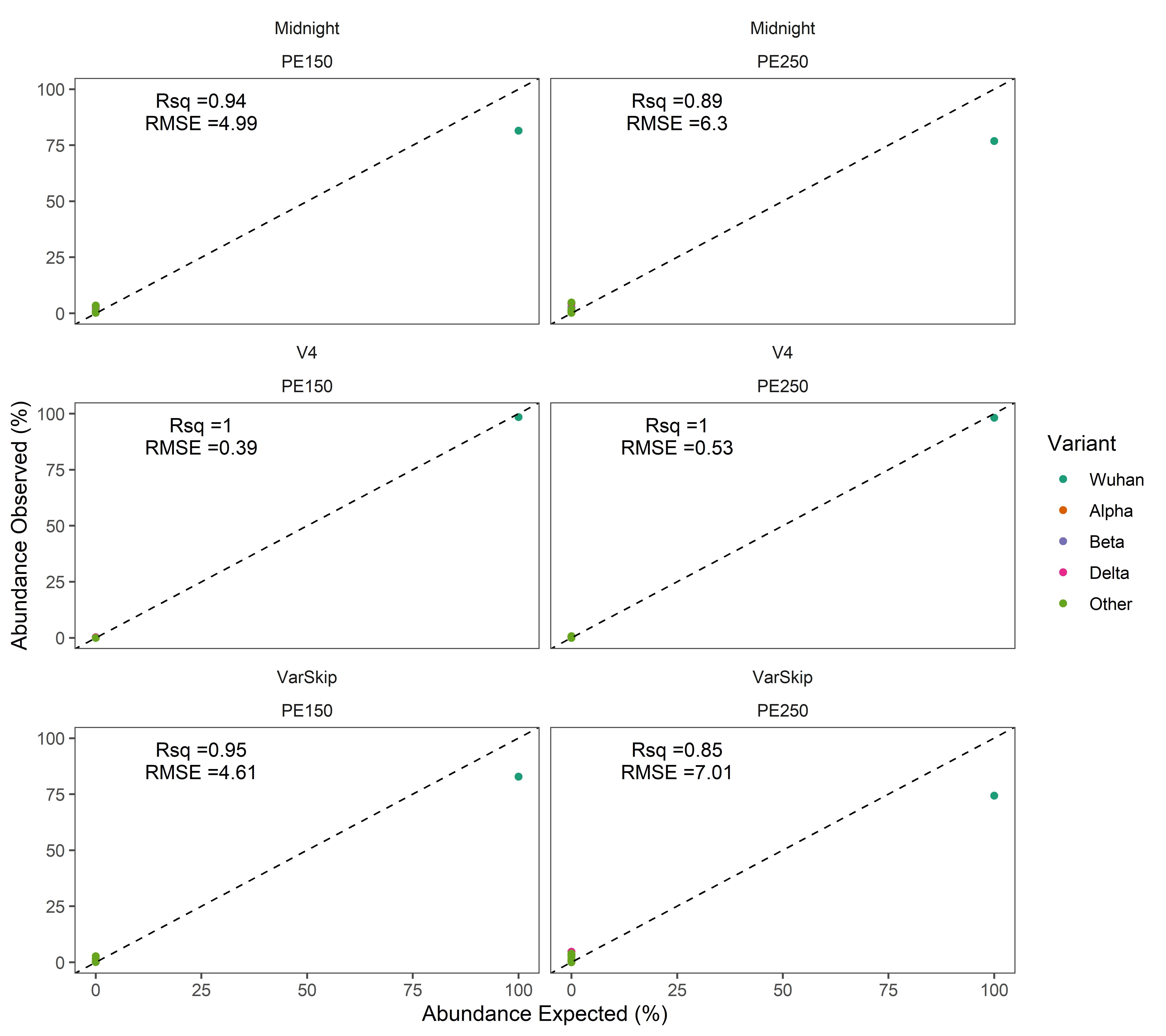

**Supplemental Figure 8. Abundance of SARS-CoV-2 variants in the SARS-CoV-2 Wuhan Twist Control Illumina Libraries, as predicted by kallisto.** The SARS-CoV-2 Wuhan Twist Control was enriched using the Freed/Midnight, ARTIC V4, and NEB VarSkip workflows, and libraries were sequenced with Illumina PE250 and PE150 reads, with approximately 10^3^ and 10^5^ reads, respectively. Sequencing reads were assessed using the kallisto workflow to estimate abundance of SARS-CoV-2 variants. Expected abundance (100% Wuhan) was compared to observed abundance, reported by kallisto. Accuracy of the abundance prediction was assessed as the R^2^ value of the one-to-one model between expected abundance and observed abundance of the SARS-CoV-2 variants in each sample.

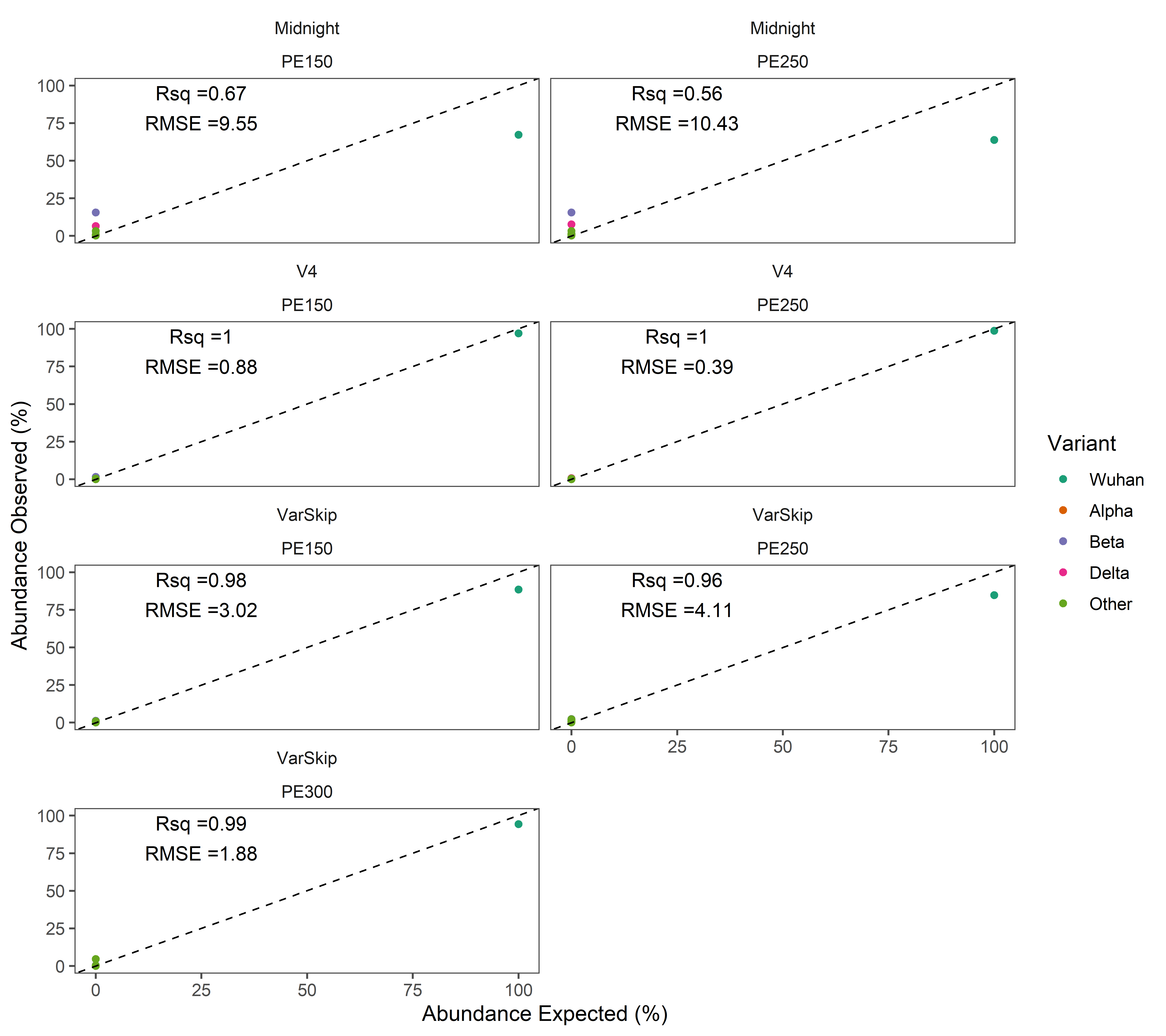

**Supplemental Figure 9. Abundance of SARS-CoV-2 variants in the heat-inactivated SARS-CoV-2 control, carried in PBS, as predicted by kallisto. The** heat-inactivated SARS-CoV-2 control, carried in PBS, was extracted using the Zymo Environmental RNA Kit. Extracted genomic material was then enriched using the Freed/Midnight, ARTIC V4, and NEB VarSkip workflows. Libraries were sequenced with Illumina PE250 and PE150 reads, with approximately 10^3^ and 10^5^ reads, respectively. Sequencing reads were assessed using the kallisto workflow to estimate abundance of SARS-CoV-2 variants. Expected abundance (100% Wuhan) was compared to observed abundance, reported by kallisto. Accuracy of the abundance prediction was assessed as the R^2^ value of the one-to-one model between expected abundance and observed abundance of the SARS-CoV-2 variants in each sample.

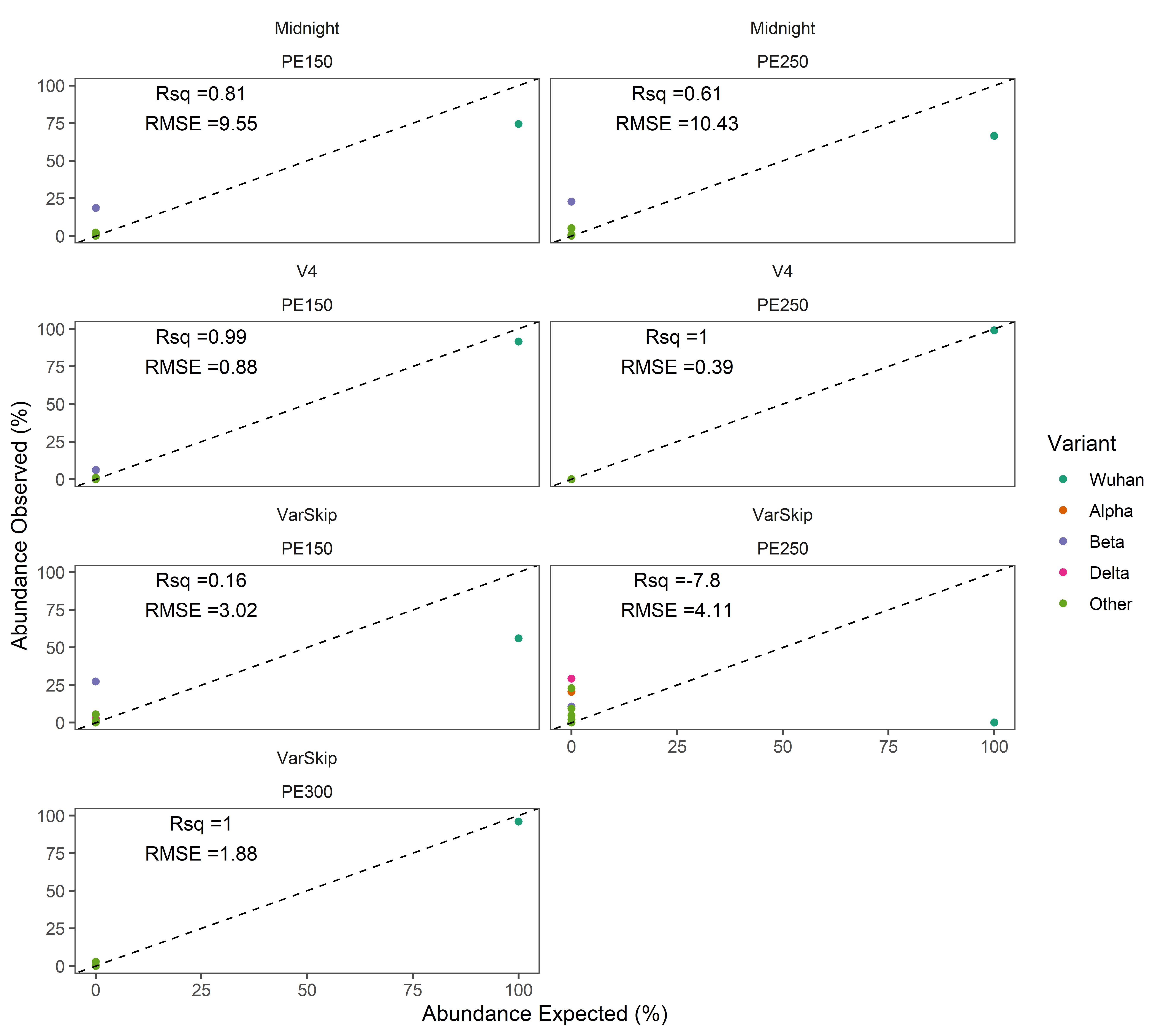

**Supplemental Figure 10. Abundance of SARS-CoV-2 variants in the heat-inactivated SARS-CoV-2 control, carried in wastewater, as predicted by kallisto.** The heat-inactivated SARS-CoV-2 control, carried in wastewater, was extracted using the Zymo Environmental RNA Kit. Extracted genomic material was then enriched using the Freed/Midnight, ARTIC V4, and NEB VarSkip workflows. Libraries were sequenced with Illumina PE250 and PE150 reads, with approximately 10^3^ and 10^5^ reads, respectively. Sequencing reads were assessed using the kallisto workflow to estimate abundance of SARS-CoV-2 variants. Expected abundance (100% Wuhan) was compared to observed abundance, reported by kallisto. Accuracy of the abundance prediction was assessed as the R^2^ value of the one-to-one model between expected abundance and observed abundance of the SARS-CoV-2 variants in each sample.

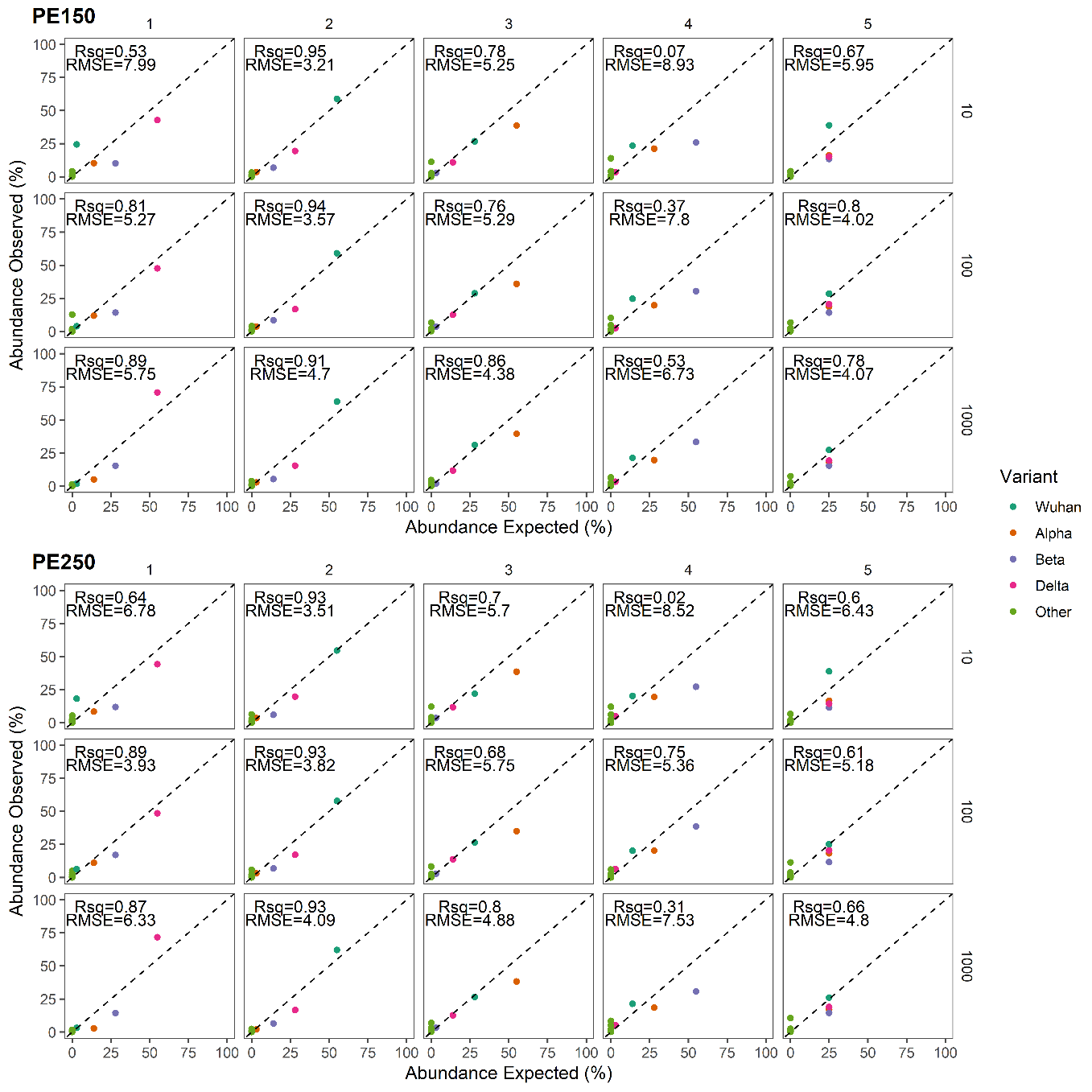

**Supplemental Figure 11. Accuracy of abundance estimates from wastewater mock communities prepared as Freed/Midnight libraries.** Synthetic genomic RNA for SARS-CoV-2 variants were combined at varying proportions to prepare five district mock communities (numbered 1 through 5), and diluted into extracted wastewater to concentrations of 10^3^, 10^2^, and 10^1^ copies μL^-1^. Mock communities were then enriched as Freed/Midnight libraries and sequenced with Illumina PE250 and PE150 chemistry, with approximately 10^3^ and 10^5^ reads, respectively. Sequencing reads were assessed using the kallisto workflow to estimate abundance of SARS-CoV-2 variants. Expected abundance, as detailed in Table 1, was compared to observed abundance, reported by kallisto. Accuracy of the abundance prediction was assessed as the R^2^ and RMSE values of the one-to-one model between expected abundance and observed abundance of the SARS-CoV-2 variants in each sample.

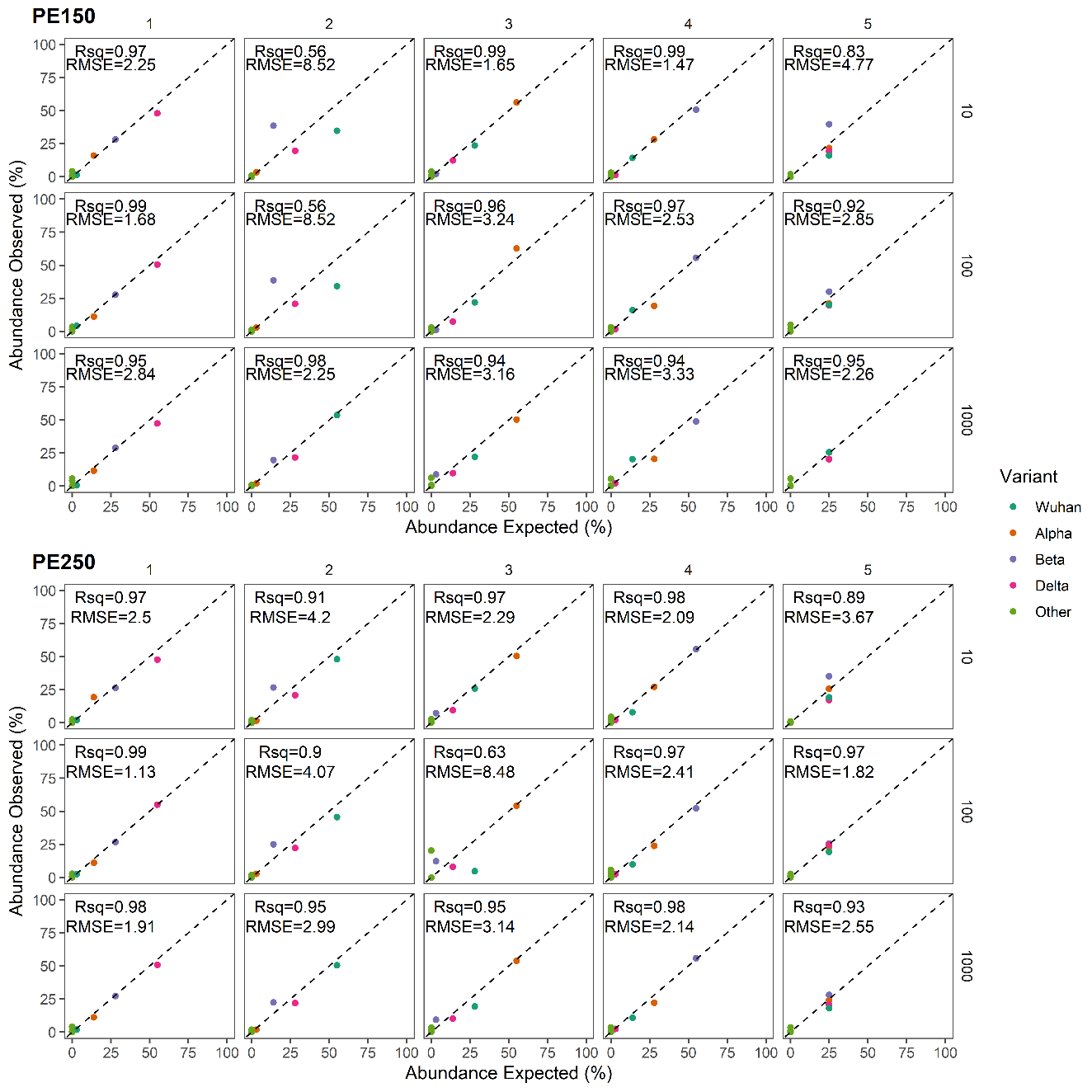

**Supplemental Figure 12.** **Accuracy of abundance estimates from wastewater mock communities prepared as ARTIC V4 libraries.** Synthetic genomic RNA for SARS-CoV-2 variants were combined at varying proportions to prepare five district mock communities (numbered 1 through 5), and diluted into extracted wastewater to concentrations of 10^3^, 10^2^, and 10^1^ copies μL^-1^. Mock communities were then enriched as ARTIC V4 libraries and sequenced with Illumina PE250 and PE150 chemistry, with approximately 10^3^ and 10^5^ reads, respectively. Sequencing reads were assessed using the kallisto workflow to estimate abundance of SARS-CoV-2 variants. Expected abundance, as detailed in Table 1, was compared to observed abundance, reported by kallisto. Accuracy of the abundance prediction was assessed as the R^2^ and RMSE values of the one-to-one model between expected abundance and observed abundance of the SARS-CoV-2 variants in each sample.

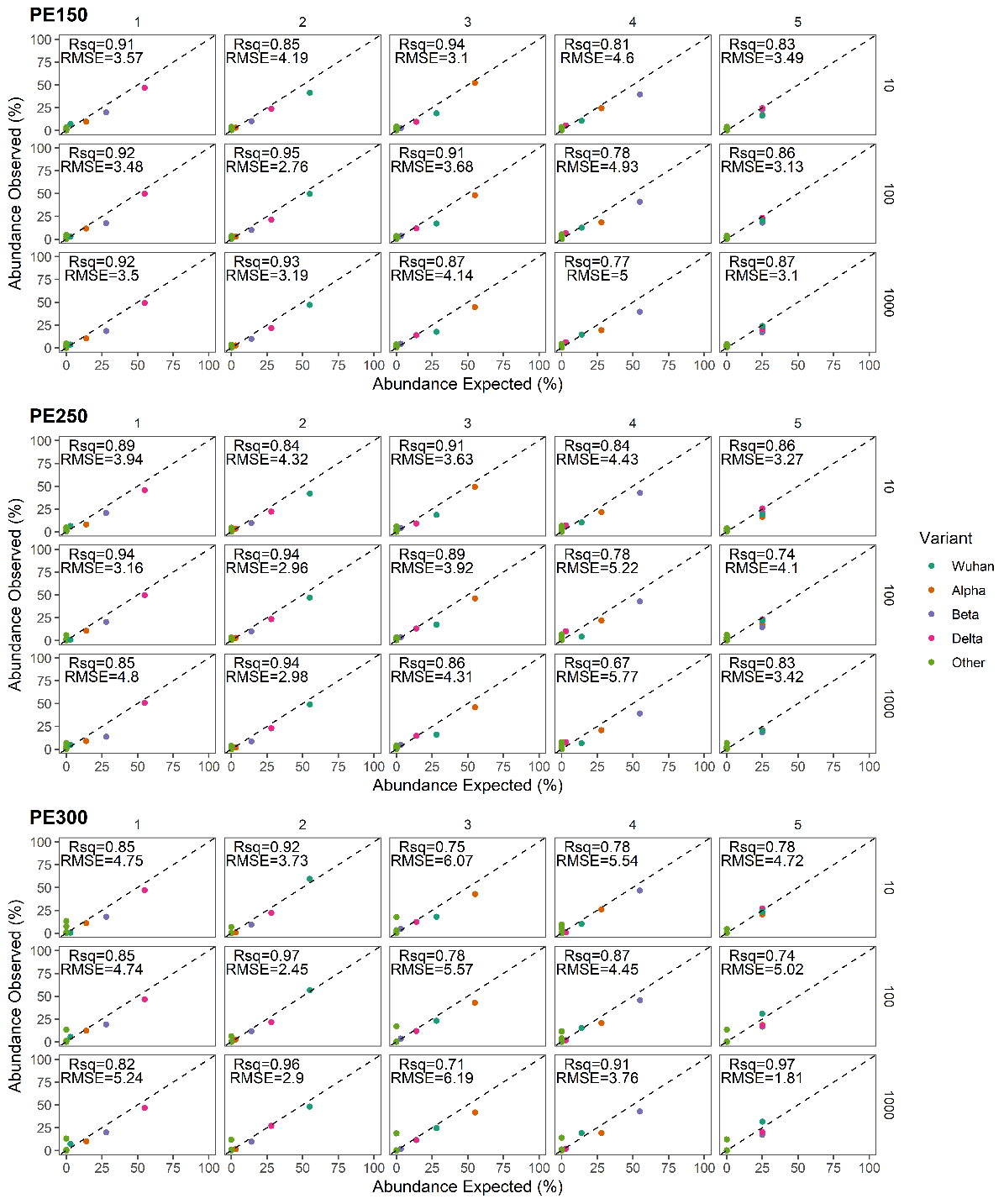

**Supplemental Figure 13.** **Accuracy of abundance estimates from wastewater mock communities prepared as NEB VarSkip libraries.** Synthetic genomic RNA for SARS-CoV-2 variants were combined at varying proportions to prepare five district mock communities (numbered 1 through 5), and diluted into extracted wastewater to concentrations of 10^3^, 10^2^, and 10^1^ copies μL^-1^. Mock communities were then enriched as NEB VarSkip libraries and sequenced with Illumina PE150, PE250, and PE300 chemistry, with approximately 10^5^ , 10^3^ , and 10^4^ reads, respectively. Sequencing reads were assessed using the kallisto workflow to estimate abundance of SARS-CoV-2 variants. Expected abundance, as detailed in Table 1, was compared to observed abundance, reported by kallisto. Accuracy of the abundance prediction was assessed as the R^2^ and RMSE values of the one-to-one model between expected abundance and observed abundance of the SARS-CoV-2 variants in each sample.

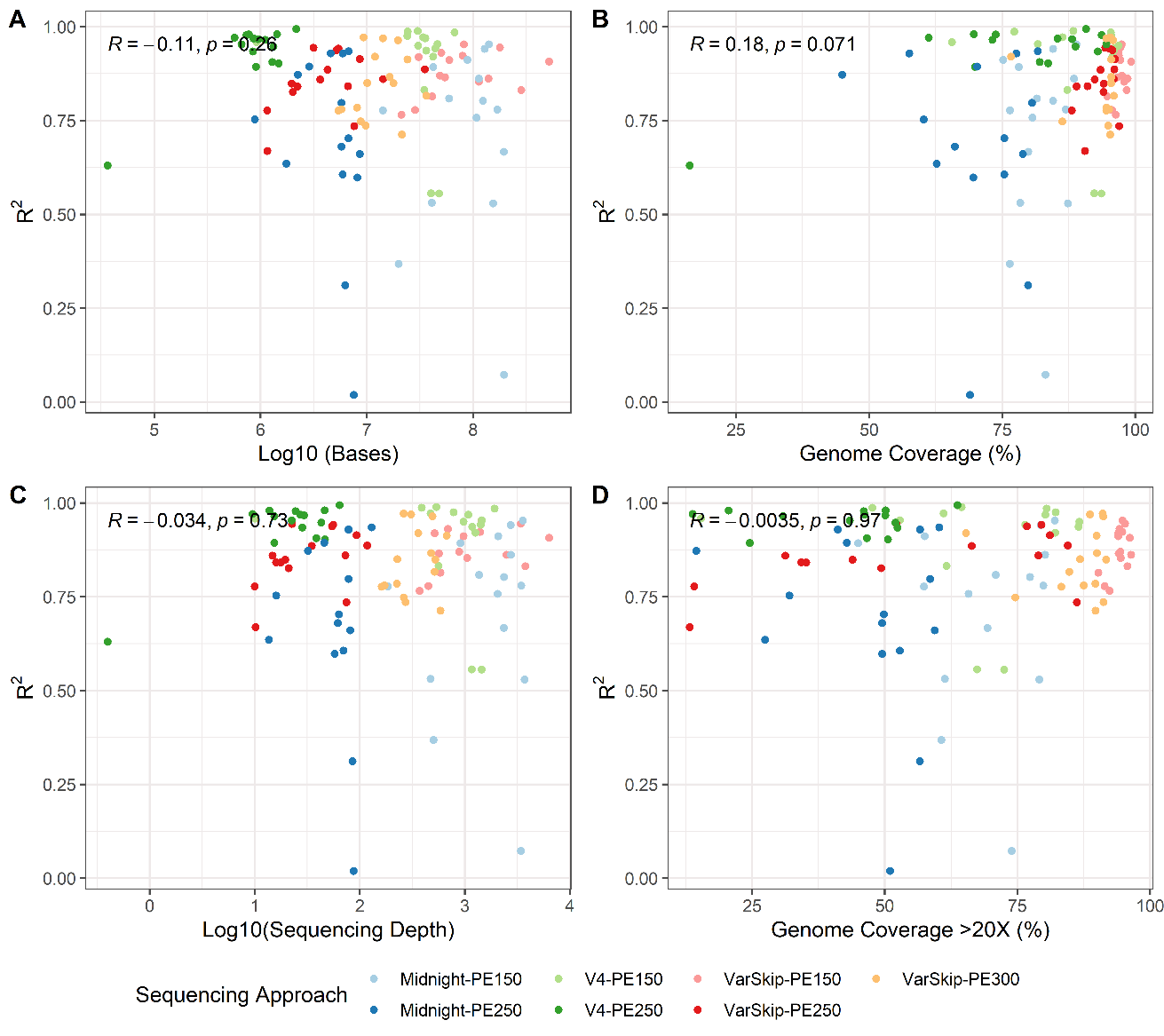

**Supplemental Figure 14.** **Variables informing the accuracy (R^2^) of variant abundance estimates from wastewater mock communities (N = 105).** Fifteen wastewater mock communities were enriched as Freed/Midnight, ARTIC V4, and NEB VarSkip libraries. Libraries were sequenced with Illumina PE250, PE150, or PE300 chemistry. Sequencing reads were assessed using the kallisto workflow to estimate abundance of SARS-CoV-2 variants. Accuracy of the abundance prediction was assessed as the RMSE value between expected observed abundances of the SARS-CoV-2 variants in each sample. The per-sample accuracy (the R^2^ value) was then assessed in correlation with the sequencing effort (A), the resulting genomic coverage (B), the average depth of genomic coverage (C), and the genomic coverage with >20X depth (D). Spearman’s Rho and p-value are reported.

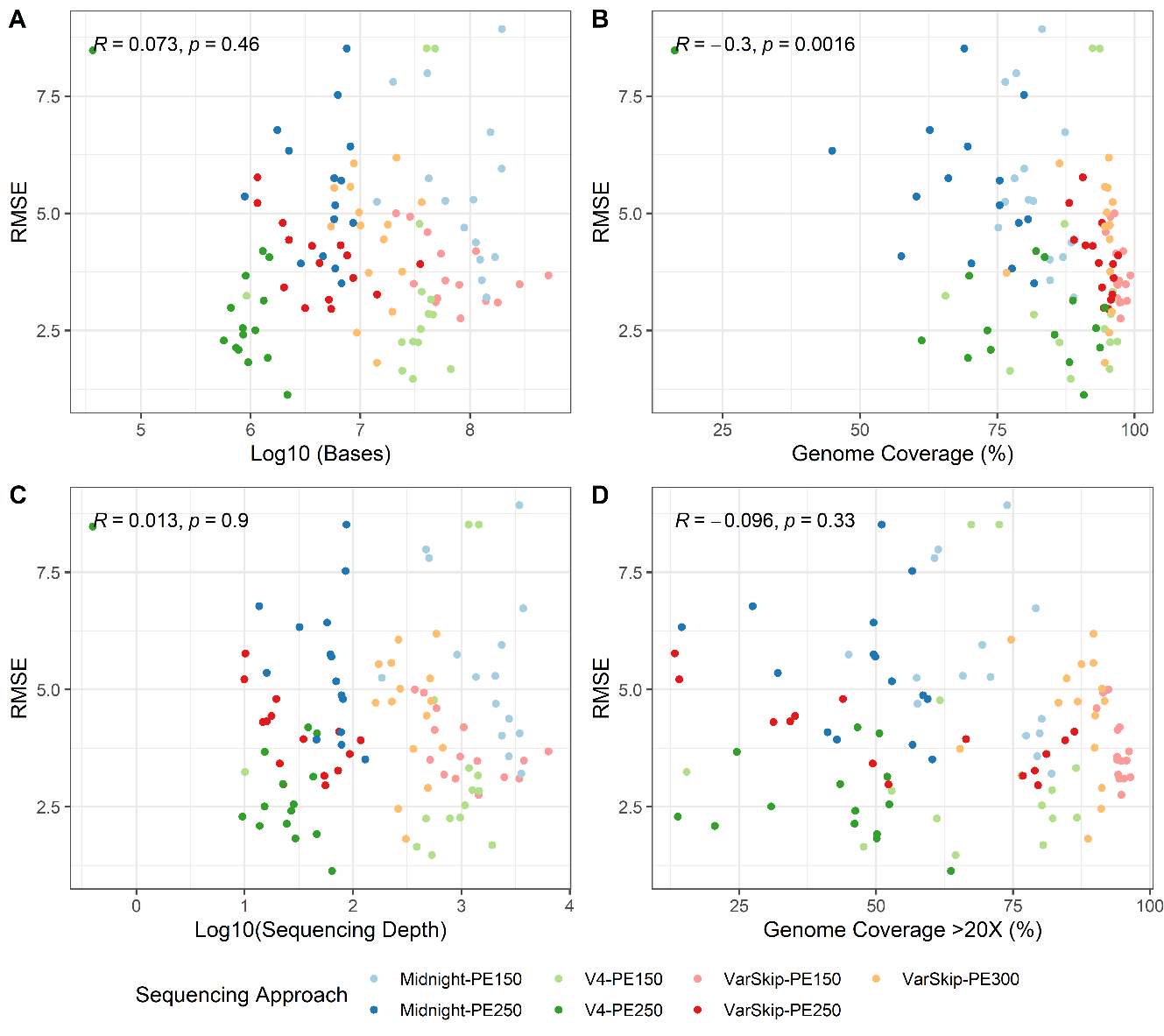

**Supplemental Figure 15.** **Variables informing the accuracy (RMSE) of variant abundance estimates from wastewater mock communities (N = 105).** Fifteen wastewater mock communities were enriched as Freed/Midnight, ARTIC V4, and NEB VarSkip libraries. Libraries were sequenced with Illumina PE250, PE150, or PE300 chemistry. Sequencing reads were assessed using the kallisto workflow to estimate abundance of SARS-CoV-2 variants. Accuracy of the abundance prediction was assessed as the RMSE value between expected observed abundances of the SARS-CoV-2 variants in each sample. The per-sample accuracy (the RMSE value) was then assessed in correlation with the sequencing effort (A), the resulting genomic coverage (B), the average depth of genomic coverage (C), and the genomic coverage with >20X depth (D). Spearman’s Rho and p-value are reported.

**Supplemental Table 2. SARS-CoV-2 Variants Spiked into Wastewater Mock Communities.** Synthetic genomic RNA for SARS-CoV-2 variants were sourced from Twist Biosciences.

| WHO Name  (Lineage) | Twist  Cat No. | Twist Control No. | GenBank/GISAID ID | GISAID Name |
| --- | --- | --- | --- | --- |
| Wuhan | 103512 | 9 | MT152824 | USA/WA2/2020 |
| Alpha  (B.1.1.7) | 103926 | 14 | EPI_ISL_710528 | England/205041766/2020 |
| Beta (B.1.351) | 104043 | 16 | EPI_ISL_678597 | South Africa/KRISP-EC-K005299/2020 |
| Delta  (AY.2) | 104539 | 29 | EPI_ISL_2693246 | USA/WA-CDC-UW21061750277/2021 |

**Supplemental Table 3. Reverse Transcription with LunaScript.**

| Component | Volume per RXN |
| --- | --- |
| LunaScript 5X MasterMix | 2 μL |
| RNA Template | 8 μL |
| TOTAL | 10 μL |
| Cycling Conditions:  1. Primer Annealing: 25°C for 2 minutes  2. cDNA Synthesis: 55°C for 10 minutes  3. Heat Inactivation: 95°C for 1 minute  4. Hold 4°C ∞ | |

**Supplemental Table 4. Amplicon Generation with Freed/Midnight Tiled Amplicon Primers.**

| Component | Volume per RXN |
| --- | --- |
| cDNA | 4.5 μL |
| NEB Q5 Hot Start 2X Master Mix | 6.25 μL |
| Primer Pool 1 or Primer Pool 2 (10 uM) | 0.55 μL |
| TOTAL | 12.5 μL |
| Cycling Conditions:  1 Initial Denaturation: 98°C for 30 seconds  2. Denaturation: 95°C for 15 seconds  3. Annealing/Extension: 65°C for 5 minutes  4. Repeat steps 2 and 3 for 35 total cycles.  5. Hold 4°C ∞ | |

**Supplemental Table 5. NEBNext Fragmentation and End Prep.**

| Component | Volume per RXN |
| --- | --- |
| Targeted cDNA Amplicons  (VarSkip and Midnight) | 13 μL |
| NEBNext Ultra II FS Enzyme Mix | 1 μL |
| NEBNext Ultra II FS Reaction Buffer | 3.5 μL |
| TOTAL | 17.5 μL |
| Cycling Conditions:  1. 37°C for 15 minutes  2. 65°C for 30 minutes  3. Hold 4°C ∞ | |

**Supplemental Table 6. Adapter ligation.**

| Component | Volume per RXN |
| --- | --- |
| End Prep Reaction Mixture | 30 μL |
| NEBNext Ultra II Ligation Master Mix | 15 μL |
| Custom iTru Stubs (5 uM) | 1.25 μL |
| TOTAL | 46.25 μL |
| Cycling Conditions:  1. 20°C for 15 minutes with heated lid off. | |

**Supplemental Table 7. iTru PCR**

| Component | Volume per RXN |
| --- | --- |
| Adapter-Ligated DNA | 7.5 μL |
| NEBNext Ultra II Q5 Master Mix | 12.5 μL |
| Custom iTru5 Primer (5 uM) | 2.5 μL |
| Custom iTru7 Primer (5 uM) | 2.5 μL |
| TOTAL | 25 μL |
| Cycling Conditions:  1. Initial Denaturation: 98°C for 30 seconds  2. Denaturation: 95°C for 15 seconds  3. Annealing/Extension: 65°C for 75 seconds  4. Repeat steps 2 and 3 for a total of 6 cycles  5. Final Extension: 65°C for 5 minutes  5. Hold 4°C ∞ | |

**Supplemental Table 8. Amplicon Generation with ARTIC V4 and NEB VarSkip Tiled Amplicon Primers.**

| Component | Volume per RXN |
| --- | --- |
| cDNA | 4.5 μL |
| NEB Q5 Hot Start 2X Master Mix | 6.25 μL |
| Primer Pool 1 or Primer Pool 2 (10 µM) | 1.75 μL |
| TOTAL | 12.5 μL |
| Cycling Conditions:  1 Initial Denaturation: 98°C for 30 seconds  2. Denaturation: 95°C for 15 seconds  3. Annealing/Extension: 63°C for 5 minutes  4. Repeat steps 2 and 3 for 35 total cycles.  5. Hold 4°C ∞ | |

**Supplemental Table 9. NEBNext End Prep (without fragmentation).**

| Component | Volume per RXN |
| --- | --- |
| Targeted cDNA Amplicons (V4) | 25 μL |
| NEBNext Ultra II End Prep Enzyme Mix | 1.5 μL |
| NEBNext Ultra II End Prep Reaction Buffer | 3.5 μL |
| TOTAL | 30 μL |
| Cycling Conditions:  1 20°C for 30 minutes  2. 65°C for 30 minutes  3. Hold 4°C ∞ | |

**Supplemental Table 9. Reference genomes for SARS-CoV-2 variants. Genomes were accessed through GISAID.**

| Strain | GISAID EIPISL | Submission Date | Country | Lineage | WHO Name |
| --- | --- | --- | --- | --- | --- |
| hCoV-19/USA/MI-MDHHS-SC39565/2022 | EPI_ISL_10266063 | 2/3/2022 | USA | BA.1 | Omicron_BA.1 |
| hCoV-19/USA/TX-TAMGHRC-SJH_129928/2022 | EPI_ISL_10271778 | 2/6/2022 | USA | BA.1 | Omicron_BA.1 |
| hCoV-19/USA/LA-BIE-LSUH002511/2021 | EPI_ISL_8479558 | 12/24/2021 | USA | BA.1 | Omicron_BA.1 |
| hCoV-19/USA/DE-Curative-177187/2022 | EPI_ISL_8976724 | 1/7/2022 | USA | BA.1 | Omicron_BA.1 |
| hCoV-19/USA/TX-TAMGHRC-L-72062-21R/2021 | EPI_ISL_9882210 | 12/20/2021 | USA | BA.1 | Omicron_BA.1 |
| hCoV-19/USA/CA-CDPH-3000325777/2022 | EPI_ISL_11908919 | 2/1/2022 | USA | BA.1 | Omicron_BA.1 |
| hCoV-19/USA/CA-CDPH-3000327137/2022 | EPI_ISL_11909111 | 2/1/2022 | USA | BA.1 | Omicron_BA.1 |
| hCoV-19/USA/CA-CDPH-3000327519/2022 | EPI_ISL_11909146 | 2/1/2022 | USA | BA.1 | Omicron_BA.1 |
| hCoV-19/USA/LA-BIE-CAT01444/2021 | EPI_ISL_15878109 | 12/23/2021 | USA | BA.1 | Omicron_BA.1 |
| hCoV-19/USA/LA-BIE-CAT01656/2021 | EPI_ISL_16461463 | 12/28/2021 | USA | BA.1 | Omicron_BA.1 |
| hCoV-19/USA/LA-BIE-LSUH002274/2021 | EPI_ISL_7799914 | 12/7/2021 | USA | BA.1.1 | Omicron_BA.1 |
| hCoV-19/USA/LA-BIE-LSUH002343/2021 | EPI_ISL_8083839 | 12/15/2021 | USA | BA.1.1 | Omicron_BA.1 |
| hCoV-19/USA/LA-BIE-LSUH002347/2021 | EPI_ISL_8083843 | 12/15/2021 | USA | BA.1.1 | Omicron_BA.1 |
| hCoV-19/USA/MS-BIE-LSUH002358/2021 | EPI_ISL_8083853 | 12/17/2021 | USA | BA.1.1 | Omicron_BA.1 |
| hCoV-19/USA/LA-BIE-LSUH002376/2021 | EPI_ISL_8083867 | 12/15/2021 | USA | BA.1.1 | Omicron_BA.1 |
| hCoV-19/USA/LA-BIE-LSUH002507/2021 | EPI_ISL_8479541 | 12/24/2021 | USA | BA.1.1 | Omicron_BA.1 |
| hCoV-19/USA/LA-BIE-LSUH002484/2021 | EPI_ISL_8479553 | 12/22/2021 | USA | BA.1.1 | Omicron_BA.1 |
| hCoV-19/USA/LA-BIE-LSUH002530/2021 | EPI_ISL_8479561 | 12/20/2021 | USA | BA.1.1 | Omicron_BA.1 |
| hCoV-19/USA/LA-BIE-LSUH002561/2021 | EPI_ISL_8687367 | 12/30/2021 | USA | BA.1.1 | Omicron_BA.1 |
| hCoV-19/USA/LA-BIE-LSUH002612/2021 | EPI_ISL_8687412 | 12/30/2021 | USA | BA.1.1 | Omicron_BA.1 |
| hCoV-19/USA/TX-TAMGHRC-L-73256-22R/2022 | EPI_ISL_10076456 | 1/21/2022 | USA | BA.2 | Omicron_BA.2 |
| hCoV-19/USA/MI-MDHHS-SC39946/2022 | EPI_ISL_10944162 | 2/20/2022 | USA | BA.2 | Omicron_BA.2 |
| hCoV-19/USA/MI-MDHHS-SC40219/2022 | EPI_ISL_12036686 | 3/31/2022 | USA | BA.2 | Omicron_BA.2 |
| hCoV-19/USA/WA-UW-22060940707/2022 | EPI_ISL_13777383 | 6/9/2022 | USA | BA.2 | Omicron_BA.2 |
| hCoV-19/USA/PA-Curative-496734/2022 | EPI_ISL_13784685 | 7/4/2022 | USA | BA.2 | Omicron_BA.2 |
| hCoV-19/USA/MI-MDHHS-SC25096/2021 | EPI_ISL_1367852 | 3/9/2021 | USA | B.1.351 | Beta |
| hCoV-19/USA/WI-WSLH-213165/2021 | EPI_ISL_1794060 | 3/26/2021 | USA | B.1.351 | Beta |
| hCoV-19/USA/OR-TRACE-LINN-033021-466/2021 | EPI_ISL_1866418 | 3/30/2021 | USA | B.1.351 | Beta |
| hCoV-19/USA/WA-UW-21042271709/2021 | EPI_ISL_2031557 | 4/22/2021 | USA | B.1.351 | Beta |
| hCoV-19/USA/CA-CDPH-3000006761/2021 | EPI_ISL_2837403 | 4/17/2021 | USA | B.1.351 | Beta |
| hCoV-19/USA/TX-TCH-TCMC03115/2021 | EPI_ISL_1621256 | 3/23/2021 | USA | P.1 | Gamma |
| hCoV-19/USA/WI-WSLH-213664/2021 | EPI_ISL_1794519 | 3/27/2021 | USA | P.1 | Gamma |
| hCoV-19/USA/CA-CDPH-3000007215/2021 | EPI_ISL_2837733 | 4/10/2021 | USA | P.1 | Gamma |
| hCoV-19/USA/NY-WMC2021-54/2021 | EPI_ISL_3046587 | 6/5/2021 | USA | P.1 | Gamma |
| hCoV-19/USA/CA-CDPH-3000130472/2021 | EPI_ISL_4511365 | 7/28/2021 | USA | P.1 | Gamma |
| hCoV-19/USA/FL-Shands-VTM-53/2021 | EPI_ISL_2989846 | 2021-01 | USA | B.1.427 | Epsilon |
| hCoV-19/USA/CA-ALSR-5090/2020 | EPI_ISL_730345 | 9/28/2020 | USA | B.1.427 | Epsilon |
| hCoV-19/USA/CA-CHLA-PLM84336997/2020 | EPI_ISL_753480 | 11/30/2020 | USA | B.1.427 | Epsilon |
| hCoV-19/USA/WA-UW-46966/2020 | EPI_ISL_756213 | 12/13/2020 | USA | B.1.427 | Epsilon |
| hCoV-19/USA/WY-WPHL-20172467/2020 | EPI_ISL_770486 | 12/30/2020 | USA | B.1.427 | Epsilon |
| hCoV-19/USA/FL-STP-Saliva-412-VAC/2021 | EPI_ISL_2989606 | 2021-02 | USA | B.1.429 | Epsilon |
| hCoV-19/USA/CA-ALSR-5112/2020 | EPI_ISL_730359 | 12/4/2020 | USA | B.1.429 | Epsilon |
| hCoV-19/USA/CA-LACPHL-AF00003/2020 | EPI_ISL_737238 | 12/18/2020 | USA | B.1.429 | Epsilon |
| hCoV-19/USA/CA-CZB-14781/2020 | EPI_ISL_738759 | 11/10/2020 | USA | B.1.429 | Epsilon |
| hCoV-19/USA/UT-UPHL-2012501214/2020 | EPI_ISL_747002 | 11/25/2020 | USA | B.1.429 | Epsilon |
| hCoV-19/USA/NY-P13D5/2021 | EPI_ISL_5156877 | 4/9/2021 | USA | B.1.525 | Eta |
| hCoV-19/USA/MI-MDHHS-SC23727/2021 | EPI_ISL_1158385 | 1/26/2021 | USA | B.1.526 | Iota |
| hCoV-19/USA/AZ-ASU4262/2021 | EPI_ISL_1743681 | 4/7/2021 | USA | B.1.526 | Iota |
| hCoV-19/USA/WI-GMF-B00437/2021 | EPI_ISL_1745217 | 4/15/2021 | USA | B.1.526 | Iota |
| hCoV-19/USA/DE-DHSS-B1084257/2021 | EPI_ISL_2107605 | 4/22/2021 | USA | B.1.526 | Iota |
| hCoV-19/USA/NY-WMC2021-15/2021 | EPI_ISL_2400450 | 4/10/2021 | USA | B.1.526 | Iota |
| hCoV-19/USA/CA-Stanford-12_S42/2021 | EPI_ISL_1379889 | 3/1/2021 | USA | B.1.617.1 | Kappa |
| hCoV-19/USA/TX-TCH-TCMC0421021/2021 | EPI_ISL_1792927 | 4/15/2021 | USA | B.1.617.1 | Kappa |
| hCoV-19/USA/CA-CZB-30484/2021 | EPI_ISL_1821470 | 3/9/2021 | USA | B.1.617.1 | Kappa |
| hCoV-19/USA/WA-UW-21041984889/2021 | EPI_ISL_2002806 | 4/19/2021 | USA | B.1.617.1 | Kappa |
| hCoV-19/USA/CA-CDPH-3000006742/2021 | EPI_ISL_2837392 | 4/15/2021 | USA | B.1.617.1 | Kappa |
| hCoV-19/USA/TX-HMH-MCoV-49453/2020 | EPI_ISL_5384223 | 7/12/2020 | USA | B.1.617.3 | Delta |
| hCoV-19/USA/FL-Path-Saliva-872/2021 | EPI_ISL_2991302 | 2021-01 | USA | P.2 | Zeta |
| hCoV-19/USA/VA-DCLS-2187/2020 | EPI_ISL_677212 | 11/12/2020 | USA | P.2 | Zeta |
| hCoV-19/USA/GA-EHC-458R/2021 | EPI_ISL_845768 | 1/5/2021 | USA | P.2 | Zeta |
| hCoV-19/USA/IL-NM5863/2021 | EPI_ISL_854460 | 1/5/2021 | USA | P.2 | Zeta |
| hCoV-19/USA/VA-DCLS-2375/2020 | EPI_ISL_925204 | 11/12/2020 | USA | P.2 | Zeta |
| hCoV-19/USA/NY-PRL-2021_03_02_00F22/2021 | EPI_ISL_1172845 | 2/28/2021 | USA | B.1.621 | Mu |
| hCoV-19/USA/TX-TCH-TCMC00443-0421/2021 | EPI_ISL_2135033 | 4/28/2021 | USA | B.1.621 | Mu |
| hCoV-19/USA/CO-CDPHE-2100938243/2021 | EPI_ISL_2309975 | 4/27/2021 | USA | B.1.621 | Mu |
| hCoV-19/USA/GA-GPHL-0610/2021 | EPI_ISL_3134240 | 7/13/2021 | USA | B.1.621 | Mu |
| hCoV-19/USA/MI-MDHHS-SC32153/2021 | EPI_ISL_3637925 | 8/9/2021 | USA | B.1.621 | Mu |
| hCoV-19/USA/CA-CZB-29195/2021 | EPI_ISL_1701259 | 3/5/2021 | USA | B.1.621.1 | Mu |
| hCoV-19/USA/LA-BIE-LSUH001328/2021 | EPI_ISL_3090983 | 7/16/2021 | USA | B.1.621.1 | Mu |
| hCoV-19/USA/TX-TCH-TCMC00652-0721/2021 | EPI_ISL_3149057 | 7/9/2021 | USA | B.1.621.1 | Mu |
| hCoV-19/USA/CA-CDPH-3000129332/2021 | EPI_ISL_4509710 | 7/27/2021 | USA | B.1.621.1 | Mu |
| hCoV-19/USA/NY-P12C1/2021 | EPI_ISL_5156854 | 4/7/2021 | USA | B.1.621.1 | Mu |
| hCoV-19/USA/WA-UW-21042164481/2021 | EPI_ISL_2031536 | 4/21/2021 | USA | B.1.617.2 | Delta |
| hCoV-19/USA/MN-UMGC-4018/2021 | EPI_ISL_2709235 | 6/7/2021 | USA | B.1.617.2 | Delta |
| hCoV-19/USA/MD-MDH-2732/2021 | EPI_ISL_2709802 | 6/9/2021 | USA | B.1.617.2 | Delta |
| hCoV-19/USA/FL-CDC-ASC210153974/2021 | EPI_ISL_2877677 | 6/24/2021 | USA | B.1.617.2 | Delta |
| hCoV-19/USA/LA-BIE-LSUH001398/2021 | EPI_ISL_3088939 | 7/13/2021 | USA | B.1.617.2 | Delta |
| hCoV-19/USA/MN-MDH-6587/2021 | EPI_ISL_2018467 | 4/24/2021 | USA | B.1.1.7 | Alpha |
| hCoV-19/USA/MN-MDH-6597/2021 | EPI_ISL_2018477 | 4/26/2021 | USA | B.1.1.7 | Alpha |
| hCoV-19/USA/NY-P13A4/2021 | EPI_ISL_5156866 | 4/9/2021 | USA | B.1.1.7 | Alpha |
| hCoV-19/USA/ND-NDDH-21667/2021 | EPI_ISL_16482911 | 11/18/2021 | USA | AY.103 | Delta |
| hCoV-19/USA/ND-NDDH-21668/2021 | EPI_ISL_16482912 | 11/30/2021 | USA | AY.83 | Delta |
| hCoV-19/USA/NC-NCSU-CORVASEQ-082521R34A7/2021 | EPI_ISL_16509625 | 8/25/2021 | USA | AY.39 | Delta |
| hCoV-19/USA/NC-NCSU-CORVASEQ-091621R24E8/2021 | EPI_ISL_16509630 | 9/16/2021 | USA | AY.103 | Delta |
| hCoV-19/USA/NC-NCSU-CORVASEQ-101921R24B11/2021 | EPI_ISL_16509636 | 10/19/2021 | USA | AY.44 | Delta |
| hCoV-19/USA/OK-OKPHLCOV0009514/2022 | EPI_ISL_15562852 | 1/3/2022 | USA | BA.1 | Omicron_BA.1 |
| hCoV-19/USA/OK-OKPHLCOV0009402/2022 | EPI_ISL_15562963 | 1/4/2022 | USA | BA.1.1 | Omicron_BA.1 |
| hCoV-19/USA/OK-OKPHLCOV0009337/2022 | EPI_ISL_15563025 | 1/6/2022 | USA | BA.1.1 | Omicron_BA.1 |
| hCoV-19/USA/OK-OKPHL0012230/2022 | EPI_ISL_15570876 | 1/25/2022 | USA | BA.1.1 | Omicron_BA.1 |
| hCoV-19/USA/AZ-ASU95961/2022 | EPI_ISL_16053693 | 1/11/2022 | USA | BA.1.1 | Omicron_BA.1 |
| hCoV-19/USA/CA-CDPH-UC2/2020 | EPI_ISL_413558 | 2/27/2020 | USA | B | Wuhan |
| hCoV-19/USA/FL_9590/2020 | EPI_ISL_424853 | 3/5/2020 | USA | B | Wuhan |
| hCoV-19/USA/VA-DCLS-0056/2020 | EPI_ISL_426464 | 4/1/2020 | USA | B | Wuhan |
| hCoV-19/USA/AZ-TG268188/2020 | EPI_ISL_426502 | 3/17/2020 | USA | B | Wuhan |
| hCoV-19/USA/NY-WCMP12B04P/2020 | EPI_ISL_427599 | 3/16/2020 | USA | B | Wuhan |
| hCoV-19/USA/CA-CDPH-6000009937/2021 | EPI_ISL_11972369 | 6/29/2021 | USA | AY.2 | Delta |
| hCoV-19/USA/CA-CDPH-N3505156/2021 | EPI_ISL_13643111 | 8/6/2021 | USA | AY.2 | Delta |
| hCoV-19/USA/CA-CDPH-N3615058/2021 | EPI_ISL_13643224 | 8/23/2021 | USA | AY.2 | Delta |
| hCoV-19/USA/TN-VUMC-SDL-SBP_8190-MS-1_0400/2021 | EPI_ISL_14590239 | 6/8/2021 | USA | AY.2 | Delta |
| hCoV-19/USA/TX-KVI27200/2021 | EPI_ISL_14721283 | 8/19/2021 | USA | AY.2 | Delta |
| hCoV-19/USA/NE-COPH-21979/2021 | EPI_ISL_17266646 | 5/26/2021 | USA | B.1.1.7 | Alpha |
| hCoV-19/USA/NE-COPH-22289/2021 | EPI_ISL_17266650 | 6/1/2021 | USA | B.1.1.7 | Alpha |
| hCoV-19/USA/CA-HLX-STM-K7FXMGFGA/2022 | EPI_ISL_15952681 | 10/29/2022 | USA | B.1.1.529 | Omicron_BA.1 |
| hCoV-19/USA/CA-HLX-STM-FUHDGYZTF/2022 | EPI_ISL_16512137 | 11/14/2022 | USA | B.1.1.529 | Omicron_BA.1 |
| hCoV-19/USA/CA-HLX-STM-U9GJCPXX7/2022 | EPI_ISL_16513051 | 11/30/2022 | USA | B.1.1.529 | Omicron_BA.1 |
| hCoV-19/USA/CA-HLX-STM-YB8VHUGYG/2022 | EPI_ISL_16603578 | 12/19/2022 | USA | B.1.1.529 | Omicron_BA.1 |
| hCoV-19/USA/CA-CDC-QDX45210011/2023 | EPI_ISL_16662441 | 1/2/2023 | USA | B.1.1.529 | Omicron_BA.1 |
| hCoV-19/USA/OK-PHL-0023373/2022 | EPI_ISL_17120797 | 2/2/2022 | USA | BA.1 | Omicron_BA.1 |
| hCoV-19/USA/SC-784AA0B9BEE/2021 | EPI_ISL_17137210 | 12/22/2021 | USA | BA.1 | Omicron_BA.1 |
| hCoV-19/USA/NY-URMC-L7RXA015-1/2022 | EPI_ISL_17239894 | 1/13/2022 | USA | BA.1 | Omicron_BA.1 |
| hCoV-19/USA/WI-OKPHL0023034/2022 | EPI_ISL_17158654 | 1/24/2022 | USA | BA.1.1 | Omicron_BA.1 |
| hCoV-19/USA/WI-SMPH-0000352/2022 | EPI_ISL_17159715 | 5/12/2022 | USA | BA.2 | Omicron_BA.2 |
| hCoV-19/USA/WI-SMPH-0000353/2022 | EPI_ISL_17159716 | 5/16/2022 | USA | BA.2 | Omicron_BA.2 |
| hCoV-19/USA/TX-HMH-M-126455/2023 | EPI_ISL_17181191 | 2/28/2023 | USA | BA.2 | Omicron_BA.2 |
| hCoV-19/USA/NV-NSPHL-23-S002879/2023 | EPI_ISL_17226821 | 3/4/2023 | USA | BA.2 | Omicron_BA.2 |
| hCoV-19/USA/CA-CDPH-7000019252/2022 | EPI_ISL_17227821 | 9/28/2022 | USA | BA.2 | Omicron_BA.2 |
| hCoV-19/USA/TX-HMH-M-115698/2022 | EPI_ISL_15089836 | 9/14/2022 | USA | BA.4 | Omicron_BA.4 |
| hCoV-19/USA/TX-HMH-MCoV-116056/2022 | EPI_ISL_15306317 | 9/20/2022 | USA | BA.4 | Omicron_BA.4 |
| hCoV-19/USA/NY-Wadsworth-22040036-01/2022 | EPI_ISL_15506298 | 8/8/2022 | USA | BA.4 | Omicron_BA.4 |
| hCoV-19/USA/TX-UTEP-B002533605/2022 | EPI_ISL_17051746 | 7/11/2022 | USA | BA.4 | Omicron_BA.4 |
| hCoV-19/USA/TX-UTEP-B002526808/2022 | EPI_ISL_17051803 | 7/18/2022 | USA | BA.4 | Omicron_BA.4 |
| hCoV-19/USA/VA-VTVAS3-GSC39965/2022 | EPI_ISL_16013473 | 10/25/2022 | USA | BA.5 | Omicron_BA.5 |
| hCoV-19/USA/NV-NSPHL-22-00089979-01/2022 | EPI_ISL_16310313 | 12/9/2022 | USA | BA.5 | Omicron_BA.5 |
| hCoV-19/USA/AR-UAMS-V05687/2022 | EPI_ISL_17109453 | 8/29/2022 | USA | BA.5 | Omicron_BA.5 |
| hCoV-19/USA/AR-V04552/2022 | EPI_ISL_17143673 | 5/27/2022 | USA | BA.5 | Omicron_BA.5 |
| hCoV-19/USA/TX-HMH-M-126413/2023 | EPI_ISL_17180729 | 2/24/2023 | USA | BA.5 | Omicron_BA.5 |
| hCoV-19/USA/MD-HP33741-PIDVYSPRHE/2020 | EPI_ISL_13491310 | 5/6/2020 | USA | B.1.503 | Beta |
| hCoV-19/USA/IL-NM-32750/2020 | EPI_ISL_13896709 | 7/17/2020 | USA | B.1.503 | Beta |
| hCoV-19/USA/NY-MSK-1774/2020 | EPI_ISL_14555516 | 5/8/2020 | USA | B.1.503 | Beta |
| hCoV-19/USA/TN-UW-5500558/2020 | EPI_ISL_2604100 | 6/3/2020 | USA | B.1.503 | Beta |
| hCoV-19/USA/CA-CZB-4857/2020 | EPI_ISL_2658572 | 9/1/2020 | USA | B.1.503 | Beta |
| hCoV-19/USA/WA-MG_LHAB49/2021 | EPI_ISL_17206541 | 7/7/2021 | USA | AY.25 | Delta |
| hCoV-19/USA/WA-MG_LHAB68/2021 | EPI_ISL_17206542 | 9/8/2021 | USA | AY.25 | Delta |
| hCoV-19/USA/WA-MG_LHAB100/2021 | EPI_ISL_17206567 | 8/15/2021 | USA | AY.25 | Delta |
| hCoV-19/USA/WA-MG_LHAB185/2021 | EPI_ISL_17212965 | 7/21/2021 | USA | AY.25 | Delta |
| hCoV-19/USA/MT-MTPHL-3957377/2021 | EPI_ISL_17260581 | 12/30/2021 | USA | AY.25 | Delta |
| hCoV-19/USA/SC-7331/2021 | EPI_ISL_17138427 | 12/25/2021 | USA | AY.39 | Delta |
| hCoV-19/USA/SC-FF16EA763IV/2021 | EPI_ISL_17141332 | 9/1/2021 | USA | AY.39 | Delta |
| hCoV-19/USA/AR-V01670/2021 | EPI_ISL_17142036 | 8/26/2021 | USA | AY.39 | Delta |
| hCoV-19/USA/WI-CDC-2-5233997/2021 | EPI_ISL_17160542 | 11/15/2021 | USA | AY.39 | Delta |
| hCoV-19/USA/WA-MG_LHAB6/2021 | EPI_ISL_17206553 | 7/29/2021 | USA | AY.44 | Delta |
| hCoV-19/USA/WA-MG_LHAB122/2021 | EPI_ISL_17206600 | 9/7/2021 | USA | AY.44 | Delta |
| hCoV-19/USA/ND-NDDH-23022/2021 | EPI_ISL_17226758 | 8/25/2021 | USA | AY.44 | Delta |
| hCoV-19/USA/NE-COPH-002080/2021 | EPI_ISL_17266558 | 12/29/2021 | USA | AY.44 | Delta |
| hCoV-19/USA/CT-Yale-17359/2021 | EPI_ISL_10654266 | 9/27/2021 | USA | AY.46.1 | Delta |
| hCoV-19/USA/NY-WMC2022-2709/2022 | EPI_ISL_13463633 | 1/4/2022 | USA | AY.46.1 | Delta |
| hCoV-19/USA/CT-Yale-7575/2021 | EPI_ISL_15573627 | 8/3/2021 | USA | AY.46.1 | Delta |
| hCoV-19/USA/CT-Yale-13738/2021 | EPI_ISL_15604332 | 11/14/2021 | USA | AY.46.1 | Delta |
| hCoV-19/USA/OK-PHL-0022092/2021 | EPI_ISL_17110603 | 12/18/2021 | USA | AY.46.1 | Delta |
| hCoV-19/USA/OK-OKPHL0004688/2021 | EPI_ISL_15567287 | 10/13/2021 | USA | AY.62 | Delta |
| hCoV-19/USA/SC-2215/2021 | EPI_ISL_17113774 | 7/6/2021 | USA | AY.62 | Delta |
| hCoV-19/USA/WV-WV123612/2021 | EPI_ISL_17117666 | 9/9/2021 | USA | AY.62 | Delta |
| hCoV-19/USA/AR-UAMS-B0439465/2021 | EPI_ISL_17126124 | 8/19/2021 | USA | AY.62 | Delta |
| hCoV-19/USA/NM-NMINBRE-2021231250/2021 | EPI_ISL_17160312 | 11/5/2021 | USA | AY.62 | Delta |
| hCoV-19/USA/WV-WV128718/2021 | EPI_ISL_17118509 | 12/27/2021 | USA | AY.75 | Delta |
| hCoV-19/USA/AR-UAMS-V02004/2021 | EPI_ISL_17121812 | 10/1/2021 | USA | AY.75 | Delta |
| hCoV-19/USA/AR-B012233/2021 | EPI_ISL_17133742 | 11/29/2021 | USA | AY.75 | Delta |
| hCoV-19/USA/NM-NMINBRE-2021122467/2021 | EPI_ISL_17160278 | 6/2/2021 | USA | AY.75 | Delta |
| hCoV-19/USA/ND-NDDH-23054/2021 | EPI_ISL_17226687 | 8/27/2021 | USA | AY.75 | Delta |
| hCoV-19/USA/WI-UW-6431/2021 | EPI_ISL_17158786 | 10/27/2021 | USA | AY.100 | Delta |
| hCoV-19/USA/NM-NMINBRE-2021237823/2021 | EPI_ISL_17160425 | 12/2/2021 | USA | AY.100 | Delta |
| hCoV-19/USA/NM-CDC-2-4678890/2021 | EPI_ISL_17160589 | 7/14/2021 | USA | AY.100 | Delta |
| hCoV-19/USA/NC-NCSU-CORVASEQ-091721R11C11/2021 | EPI_ISL_17179386 | 9/17/2021 | USA | AY.100 | Delta |
| hCoV-19/USA/ND-NDDH-22989/2021 | EPI_ISL_17226653 | 11/16/2021 | USA | AY.100 | Delta |
| hCoV-19/USA/NE-CDC-2-4679188/2021 | EPI_ISL_17160579 | 7/20/2021 | USA | AY.103 | Delta |
| hCoV-19/USA/ND-NDDH-23023/2021 | EPI_ISL_17226759 | 8/25/2021 | USA | AY.103 | Delta |
| hCoV-19/USA/NE-COPH-000916/2021 | EPI_ISL_17266555 | 9/13/2021 | USA | AY.103 | Delta |
| hCoV-19/USA/OK-PHL-0026055/2021 | EPI_ISL_17126535 | 11/29/2021 | USA | AY.118 | Delta |
| hCoV-19/USA/AR-B011819/2021 | EPI_ISL_17133490 | 10/29/2021 | USA | AY.118 | Delta |
| hCoV-19/USA/SC-7126/2021 | EPI_ISL_17138253 | 12/22/2021 | USA | AY.118 | Delta |
| hCoV-19/USA/NC-NCSU-CORVASEQ-091721R11E8/2021 | EPI_ISL_17179384 | 9/17/2021 | USA | AY.118 | Delta |
| hCoV-19/USA/ND-NDDH-23025/2021 | EPI_ISL_17226761 | 8/26/2021 | USA | AY.118 | Delta |
| hCoV-19/USA/SC-7158/2021 | EPI_ISL_17138284 | 12/22/2021 | USA | AY.119 | Delta |
| hCoV-19/USA/WV-WV129463/2022 | EPI_ISL_17145320 | 1/15/2022 | USA | AY.119 | Delta |
| hCoV-19/USA/WA-MG_LHAB8/2021 | EPI_ISL_17206569 | 7/29/2021 | USA | AY.119 | Delta |
| hCoV-19/USA/ND-NDDH-23014/2021 | EPI_ISL_17226750 | 8/24/2021 | USA | AY.119 | Delta |
| hCoV-19/USA/NE-COPH-001200/2021 | EPI_ISL_17266556 | 11/29/2021 | USA | AY.119 | Delta |

**Supplemental Text. Details of Statistical Analyses.**

Normality of continuous data was assessed using the Shapiro–Wilk test in the R stats package with default parameters. Non-parametric data were compared across groups using the Kruskal-Wallis test, followed by the Dunn pairwise post-hoc test with Holm-adjusted p-values. The Kruskal-Wallis and Dunn tests were conducted using the rstatix package with default parameters. Estimates of variant abundance were reported by kallisto. The accuracy of the abundance prediction was assessed as the root-mean-squared error and the R^2^ value between expected observed abundances of the SARS-CoV-2 variants in each sample.
